# Virulence shift in Type X *Toxoplasma gondii*: natural cross QTL identifies ROP33 as rodent Vir locus

**DOI:** 10.1101/2021.03.31.437793

**Authors:** A Kennard, MA Miller, A Khan, M Quinones, N Miller, N Sundar, ER James, K Greenwald, DS Roos, ME Grigg

## Abstract

How virulent parasites are maintained in nature is an important paradigm of eukaryotic pathogenesis. Here we used population genetics and molecular methods to study the evolution and emergence of genetic variants of the protozoan parasite *Toxoplasma gondii,* referred collectively as Type X (HG12), recovered from a threatened marine mammal species. Specifically, 53 *T. gondii* strains were isolated from southern sea otters (SSO) that stranded between 1998-2004 with *T. gondii* infection (ranging from chronic incidental infections to fatal encephalitis). Over 74% of these SSO, collected throughout their geographic range, were infected with Type X, based on multi-locus PCR-DNA sequencing. Depending on the locus investigated, Type X strains possessed one of three allelic types that had independently assorted across the strains examined; either genetically distinct alleles, referred to as “γ” or “8”, or a Type II allele. Phylogenetic incongruence among locus-specific trees, genome-wide CGH array and WGS analyses confirmed that Type X is a sexual clade of natural recombinants that resemble F1 progeny from a genetic cross between Type II and a mosaic of two distinct “γ” or “δ” ancestries. A single Type X genotype (19/53; 36%) that largely caused subclinical chronic infections in SSO, was highly pathogenic to mice (LD_100_= 1 parasite). To determine whether murine virulence genes could be mapped within this population of natural isolates, we performed a genome scan and identified four QTLs with LOD scores greater than 4.0. Targeted disruption of ROP33, the strongest candidate from among 16 genes within the highest QTL on Chromosome VIIa established ROP33 as a murine virulence locus. The ability of this highly pathogenic mouse-virulent *T. gondii* clone to expand its environmental niche and infect a majority of SSO supports a virulence shift model whereby generalist pathogens like *T. gondii* utilize their sexual cycles to produce new strains that possess an expanded biological potential. Such a trait enables pathogens to extend their host range or be naturally selected within their vast intermediate host range to maximize transmission. Our work establishes a rationale for how virulent strains can be maintained cryptically in nature across a pathogen’s broad host range, and act as potential reservoirs for epidemic disease.

**Importance:** Waterborne outbreaks of protozoal parasites are capable of causing fatal disease in a wide range of animals, including humans. Population expansion of felids in addition to anthropogenic changes near marine estuarine environments may facilitate marine wildlife exposure to highly infectious *Toxoplasma gondii* oocysts shed in felid feces. Infected cats shed millions of environmentally-resistant *T. gondii* oocysts that can be widely dispersed by storm events. In North America *T. gondii* is thought to possess a highly clonal population structure dominated by 4 clonal lineages (I, II, III, and X). Population genetic analysis of 53 *T. gondii* isolates collected longitudinally from SSO infected with *T. gondii* that stranded between 1998-2004 identified 74% of otters infected with Type X *T. gondii*, and that Type X is not a clonal lineage, but rather a recombinant clade of at least 12 distinct strains consistent with a recent genetic cross. Importantly, one Type X haplotype was isolated from 36% of southern sea otters (*Enhydra lutris neries*) across their geographic range in California. This haplotype was highly pathogenic to mice but caused relatively benign infections in SSO. A genome scan was performed to identify a virulence locus; a secreted serine threonine kinase (ROP33) that enhanced pathogenicity in laboratory mice, but not sea otters. Our data support a virulence shift model whereby generalist pathogens like *T. gondii* utilize their sexual cycles to produce virulent strains that can be maintained cryptically in nature, according to their differential capacity to cause disease within the pathogen’s broad intermediate host range. This type of “host selection” has important public health implications. Strains capable of causing fatal infections can persist in nature by circulating as chronic infections in resistant intermediate host species that act as reservoirs for epidemic disease.

## Introduction

How successful microbes maintain reservoirs of virulent strains in nature is understudied and is an important paradigm of infectious disease. Whereas many studies document how microbial pathogens acquire or admix genetic material to influence their pathogenicity, relatively less is known how emergent strains of varying pathogenicity are selectively maintained in nature. Viruses can undergo reassortment to rapidly produce admixture lines that possess a range of altered biological potential, including virulence [1]. Bacteria can transfer mobile plasmid elements that encode pathogenicity islands via conjugation, conferring an altered virulence potential to recipients [2–4]. Among fungi and protozoa, genomic variation can be generated by genetic hybridization and produce novel admixture lines capable of causing disease outbreaks or altering host range [5–7]. Less clear are the mechanisms that allow highly prevalent pathogens to maintain virulent strains cryptically in nature. We sought to test whether generalist pathogens, such as the protozoan parasite *T. gondii* can leverage its broad range of intermediate hosts to selectively partition parasite genotypic diversity and virulence potential.

*Toxoplasma gondii* is a highly successful and prevalent protozoan pathogen that infects diverse wildlife, livestock, and 20-80% of humans worldwide [8–10]. This parasite’s success in nature is largely attributed to its highly flexible life cycle. It is propagated asexually after ingestion of tissue cysts or sporulated oocysts among essentially all warm-blooded vertebrates. It can also propagate sexually within its definitive felid host by self-mating (when a single genotype undergoes fertilization and sexually clones itself) or by out-crossing (producing as many as 10^8^ genetic hybrids that are transmissible as environmentally stable, highly infectious oocysts). Genetic hybridization has been shown to produce genotypes that possess a wide spectrum of biological potential, including altered virulence or a capacity to expand as successful epidemic clones [7, 11, 12]. Importantly, the broad spectrum of disease states caused by these admixture clones is highly dependent on both parasite genetics and host animal species [8, 13–15]. In rodents, low dose inocula of recombinant Type I *T. gondii* strains are uniformly lethal to laboratory mice [16], whereas Type II strains are considerably less virulent and routinely establish chronic infections that are transmissible to other hosts [14]. In contrast, wild-derived CIM mice possess resistant alleles of immunity-related GTPases (IRGs), which inactivate parasite virulence-associated secreted kinases (ROP18/ROP5) and render mice resistant to *T. gondii* infection, including virulent Type I strains [17, 18]. Likewise, some laboratory rat strains and species are resistant to Type I infections and fail to transmit parasites [19]. The molecular basis for rat resistance is an allele-dependent activation of the NLRP1 inflammasome that results in macrophage pyroptosis and inhibition of parasite growth in Lewis (LEW) and Sprague Dawley (SD) rats, but not Brown Norway (BN) or Fischer CDF rats, which go on to establish chronic, transmissible infections of mouse-virulent Type I strains [20–22]. The result of this host-parasite genetic interplay determines the potential for a disease-producing virulent clone to cause either a non-transmissible infection or be maintained cryptically in a strain-specific, or host species-specific manner. While laboratory studies suggest that virulent strains can be maintained cryptically across the pathogen’s host range [23], this phenomenon has not been fully investigated, nor has it been demonstrated to occur in a natural setting.

Between 1998 and 2004, *T. gondii* was reported as a significant cause of morbidity and mortality in southern sea otters (*Enhydra lutris nereis*), a federally listed threatened species [24, 25]. Detailed necropsy and histopathology studies identified a wide spectrum of outcomes among *T. gondii*-infected sea otters, ranging from chronic asymptomatic infections to fatal systemic disease [25, 26]. Prior studies also suggested that most southern sea otters were infected by a single outbreak clone, referred to as Type X (or HG12) [25]. Based on multi-locus PCR-DNA sequencing (MLST) using a limited set of genotyping markers, Type X was defined as the fourth clonal lineage in the US [27] and was highly prevalent in sylvatic niches, representing 47% of chronic, subclinical, or mild *T. gondii* infections in examined US wildlife [28, 29]. However, the broad range of post-infection outcomes observed in stranded southern sea otters was not consistent with infection by a single clone. Other possibilities proposed to explain this variation in disease susceptibility included co-infection [30], and exposure to immune-suppressive environmental pollutants, or toxins [31]. More recently, whole genome sequencing (WGS) performed on 62 globally-distributed *T. gondii* isolates suggested that MLST analyses may not be sufficiently resolved to predict clonotypes because they survey only limited genetic heterogeneity [15]. Although a previous study concluded that Type X was comprised of two genotypes [32], these two genotypes differed by only a few SNPs, so it was not clear whether strain variation was contributing to sea otter disease outcome in this novel host-parasite interaction.

To further investigate the extent of genetic diversity among naturally circulating Type X isolates, we sequenced a wide selection of linked and unlinked markers across the nuclear and organellar genomes for 53 *T. gondii* isolates collected longitudinally from southern sea otters that stranded in California, USA from 1998 through 2004. Based on initial genotyping results, we selected 14 sequence-confirmed Type X isolates for WGS. Analysis of our data support a model whereby Type X exists as a recombinant clade of strains that resemble F1 progeny from a natural cross between a Type II strain and a novel genotype that has not been identified previously in nature. We show that one Type X genotype was widely distributed in southern sea otters and caused the majority of *T. gondii* infections, which were largely subclinical in sea otters. This same genotype was highly pathogenic to laboratory mice, and a population level, forward genetic approach using WGS mapped a mouse virulence gene to a novel parasite-specific secreted kinase ROP33, also referred to as WNG3 [33]. Our data suggest that *T. gondii* can leverage its broad range of intermediate host species to partition its genetic diversity such that infected hosts play a central role in the selection, expansion, and maintenance of cryptically virulent *T. gondii* strains. In effect, these infected hosts act as reservoirs that have the potential to cause epizootics or epidemics in other host species.

## Results

### Genetic characterization and pathogenesis of *T. gondii* strains isolated from southern sea otters

During the sample period (1998-2004), pathological examinations carried out on southern sea otters that stranded along the California coast identified numerous systemic protozoal infections caused by *T. gondii, Sarcocystis neurona*, or co-infection with both parasites [25, 29, 32, 34]. A primary goal of this study was to determine if *T. gondii*-associated sea otter mortality was dependent, at least in part, by the protozoal genotype causing infection. In total, 53 *T. gondii* isolates collected longitudinally from 1998 through 2004 from minimally decomposed, stranded southern sea otters were included in this study. Isolates were obtained throughout the sea otter range along the central California coast (Supplemental Figure 1). For nearly all isolates (n=52), sea otter IFAT titers, stranding locations, sex, and *S. neurona* co-infection status was available. Among 53 sea otters examined, *T. gondii* was considered the primary cause of death among 8 animals (15%), a contributing cause of death among 11 sea otters (21%) and determined to have caused incidental, relatively benign infections in the remaining 34 sea otters (64%) (Supplemental Figure 1).

Initial genetic analysis performed on a subset of 35 sea otter *T. gondii* isolates established that 21 were a new genetic type, referred to as Type X (also known as Haplogroup 12, or HG12), but this conclusion was limited to sequence typing at a single locus, GRA6 [25]. Hence, we developed a multi-locus (MLST) PCR-DNA sequence-based typing scheme using 4 additional unlinked markers (BSR4, BAG1, SAG3, and ROP1) that possess a wide-range of phylogenetic strength, including markers under neutral (BSR4), diversifying (GRA6, SAG3, ROP1), and purifying (BAG1) selection to determine the *T. gondii* genotypes that caused infection among 53 sea otter *T. gondii* isolates. Typing solely at the GRA6 marker identified three sequence types, either a Type II allele in 16 (30%) isolates, or two non-archetypal alleles, previously identified as X or A [25, 32], in 25 (47%) and 12 (23%) isolates, respectively. Expanding the genetic analysis across the five markers showed that 2 of 16 isolates with a Type II allele at GRA6 were predicted to be non-Type II recombinant strains, because they possessed non-archetypal alleles at ROP1 (sea otter 3675) or BSR4 and BAG1 (sea otter 3671) (Figure 1A).

**Figure 1:**
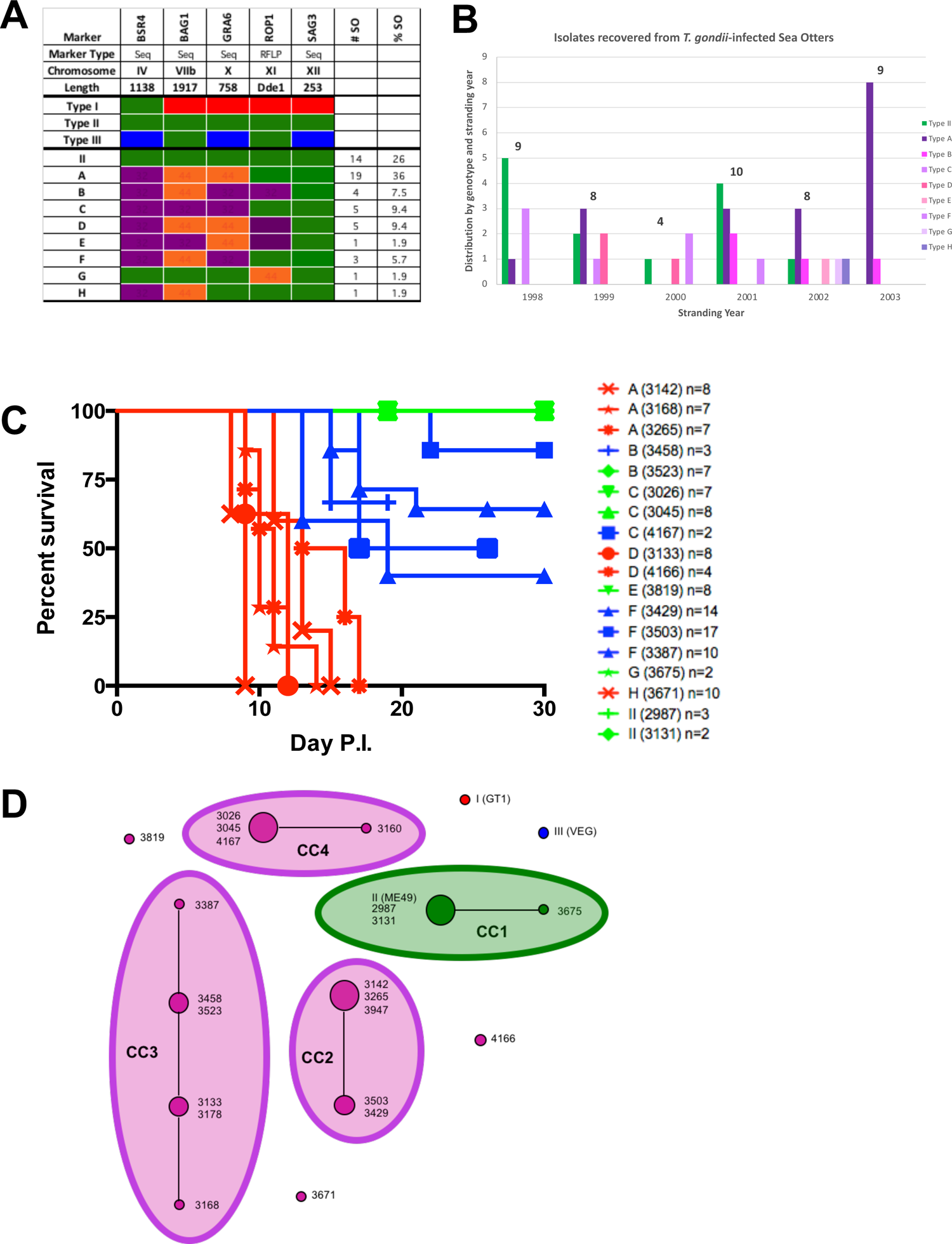
Parasite diversity and mouse virulence within the *Toxoplasma gondii* Type X clade infecting southern sea otters (*Enhydra lutris nereis*) A) Fifty-three *T. gondii* isolates from southern sea otters necropsied 1998-2004 were genotyped using 5 unlinked markers, resulting in the identification of 9 distinct *T. gondii* genotypes. Type I, II, and III alleles are represented by red, green, and blue bars, respectively. Sea otters were infected with *T. gondii* strains that possessed either non-archetypal alleles (purple or orange bars) or a Type II allele (green bars) at the loci examined. For each distinct genotype, the number of *T. gondii* infected sea otters (#SO) and its percentage of total infections (%SO) is shown. Each distinct genotype was comprised of the following *T. gondii* isolates which corresponds to the sequential stranding number of each stranded sea otter: **A** (3097, 3142, 3168, 3265, 3483, 3488, 3520, 3637, 3659, 3744, 3786, 3821, 3865, 3947, 3950, 4003, 4045, 4071, 4151), **B** (3458, 3523, 3728, 3897), **C** (3026, 3045, 3077, 3160, 4167), **D** (3133, 3178, 3183, 3451, 4166), **E** (3819), **F** (3387, 3429, 3503), **G** (3675), **H** (3671), **II** (2987, 2994, 3005, 3009, 3087, 3131, 3208, 3396, 3521, 3576, 3587, 3636, 3739, 4181). B) Distribution of isolates recovered from *T. gondii-*infected sea otters by genotype and stranding year (1998-2003): Type II (green) versus Type X haplotype (various shades of purple). Although the sample size was small, Type II infections appeared to cluster temporally, whereas Type X haplotype A infections (dark purple bars), isolated from sea otters with predominantly subclinical or incidental infections, appeared to increase in the final year of the study. C) Cohorts of CD1 mice were infected with 50 tachyzoites each intraperitoneally with various *T. gondii* genotypes isolated from southern sea otters. Each mouse cohort was infected with one of 18 isolates representing all singletons (**E, G, H**), and at least two isolates each from the remaining 6 distinct genotypes (**II, A, B, C, D, F**). Mouse seroconversion and survival was monitored for 30 days. At least two independent infection experiments were performed using 5 mice for each isolate; results shown are only for mice that seroconverted and/or died acutely during infection. Strains are grouped and colored based on their genotype and virulence in mice. **Red**; mouse virulent, **Blue**; intermediate mouse virulent (some mice survived acute infection); **Green**; mouse avirulent (all mice survived acute infection). The number of mice infected by each genotype (as indicated by acute death and/or seropositivity) and their relative virulence is indicated at right. D) eBURST analysis to determine linkage disequilibrium across 17 nuclear-encoded linked and unlinked markers identified 4 clonal complexes and 3 unique Type X genotypes among 21 isolates selected from the 9 distinct clades in Figure 1A. Isolate colors represent the lineage Type from A: red (Type I), green (Type II), blue (Type III), and purple (Type X; all haplotypes (A-H)). Dot size is proportional to the number of isolates with that genotype; larger dots are multi-isolate genotypes. Clonal complexes (CC) are indicated by lines connecting isolates and are highlighted with ovals corresponding to their allelic identity (X-purple, II-green)

### Possible point-source exposure of Type II *T. gondii* strains in southern sea otters

In total, fourteen otters were infected by Type II strains because they possessed only Type II alleles at all 5 assessed loci, including 12 (86%) otters that stranded within a 28 km span from Moss Landing to Monterey Bay; this discreet clustering of Type II *T. gondii*-infected sea otters is suggestive of regional or point source exposure. The other two Type II-infected otters were recovered from Pismo Beach, approximately 282 km south of the potential region of point source exposure. Although most Type II-infected otters (9/14; 64%) were collected during the first half of the study (Supplemental Figure 1B), due to the small sample size, this could be due to random chance. In our sample of 14 Type II *T. gondii* strains, more male sea otters were infected than females (11 vs. 3, respectively), and males were shown to be at significantly higher risk for Type II *T. gondii* infection, when compared with all other strains via a Fisher’s two-sided exact test (p=0.028). In mice, Type II strains are considered subclinical or avirulent because they require high dose infections, immunosuppression, or mutations that affect host immune signaling pathways in order to cause mortality [14]. In sea otters, infection with Type II strains was generally incidental, with only 1/14 (7%) sea otters succumbing to *T. gondii* infection as a primary cause of death (Supplemental Figure 1).

### Type X is a recombinant clade of strains and pathogenicity is dependent on haplotype

Of the remaining 39 (74%) *T. gondii* isolates, a maximum of just three alleles was identified at the 5 sequenced loci: either a canonical Type II allele or one of two genetically distinct alleles, referred to as “γ” or “δ”, that appeared to segregate independently across the isolates (Figure 1A). In total 8 distinct haplotypes, designated A-H, were resolved based on their differential inheritance of the limited alleles, consistent with recombination between a Type II strain and a mosaic of two distinct ancestries as the most plausible explanation for the genetic relationship among the non-Type II strains. The Type X haplotype designated A was most common (19/39; 49%); was widely distributed across the entire southern sea otter range (Supplemental Figure 1), and was isolated from sea otters stranding every year except 2000. Infection with haplotype A, in similar to Type II infection, was likewise generally incidental, with only 1/19 (5%) haplotype A sea otters succumbing to *T. gondii* infection as the primary cause of death (Supplemental Figure 1). Another sea otter infected with haplotype A *T. gondii* that died acutely was co-infected with *Sarcocystis neurona*, and it was not possible to differentiate the relative contribution between both parasites in relation to the cause of death. In addition, although our sample size was small and thus not definitive, most of the Type X haplotype A isolates (14/19) were collected in the last 3 years of the study, which may suggest that this genotype expanded relative to other genotypes across the seven year study. In addition, Type X haplotype A strains represented the majority of sea otter *T. gondii* infections that yielded isolates (8/9) in 2003 (Figure 1B). Conversely, infection of sea otters with *T. gondii* Type X haplotype F (3/3) was highly pathogenic; these otters were significantly more at risk from fatal toxoplasmosis, compared to otters infected with haplotype A (Fisher’s two-sided exact test [p=0.036])

### Acute virulence in mice is dependent on Type X haplotype

Linking mortality in sea otters to *T. gondii* genotype in a wild animal population is particularly challenging given the many confounding variables that may influence disease outcome, including exposure to chemical pollutants, immunosuppression, or co-infection with other pathogens such as *S. neurona*. In contrast, studies using peritoneal infection models of *T. gondii* in laboratory mice have been instrumental in identifying causal linkages between *T. gondii* genotype and virulence. Using an inoculum of 50 parasites, mouse avirulent strains (*e.g.,* Type II) establish chronic, transmissible infections, whereas mouse virulent strains (*e.g.,* Type I) die acutely within 10-14 days [7].

Among the sea otter *T. gondii* isolates, three mouse virulence phenotypes were identified: virulent (red; all mice died acutely), intermediate virulent (blue; some mice survived acute disease), and avirulent (green; all mice survived acute infection and became chronically infected). The two otter isolates that possessed a Type II MLST (2987, 3131) were avirulent (Figure 1C; green) and phenocopied the infection kinetics of two well studied Type II lines; Me49 and 76K. Among the Type X haplotypes, E and G were also avirulent. Three otter isolates from haplotype A, the genotype associated with most infections in sampled sea otters, were highly pathogenic to mice; all infected mice died acutely (Figure 1C; red). Haplotypes D and H were likewise highly virulent in mice. The three haplotype F otter isolates all possessed an intermediate virulence phenotype (Figure 1C; blue). Interestingly, the virulence phenotypes for the 5 isolates from haplotypes B and C were mixed, either avirulent (green) or intermediate virulent (blue) (Figure 1C). Because the Type X isolates displayed a range of virulence phenotypes, our data suggest that the Type X group represents a clade of recombinants from a genetic admixture of a mouse avirulent Type II strain and at least one other parent that is mouse virulent, rather than expansion of a single clonal lineage. In addition, not all isolates within each subgroup displayed equivalent virulence kinetics; isolates from the B and C haplotypes were either avirulent or possessed intermediate virulence, suggesting that the 5 marker MLST was insufficiently resolved to capture the full range of genetic and phenotypic diversity of *T. gondii* isolates infecting SSO. To explore this possibility, 21 isolates (18 isolates assayed through mice plus 3 additional isolates, one each from haplotypes A, C and D), were further genotyped using 18 genetic markers (17 nuclear-encoded single copy gene loci plus one microsatellite marker), representing an expanded set of linked and unlinked loci across the genome. Two additional markers, one each from the organellar genomes of the apicoplast and mitochondria, were also included to determine the ancestry of each maternally inherited genome. This represented a significant increase in the number of markers (n = 20), when compared with those used to conclude that Type X (HG12) is a clonal lineage [10, 27, 35]. This 20 marker MLST method provided 15,404 bp of sequence information to determine the genetic relationship among *T. gondii* strains isolated from SSO (Supplemental Table 1).

### Type X is comprised of 12 distinct haplotypes by an expanded MLST analysis

To establish the number of haplotypes among the 21 isolates described in the previous section, sequence data for the 17 nuclear-encoded single copy gene loci was concatenated and analyzed using eBURST (Figure 1D). eBURST resolves strains into clonal complexes (CC) that are in linkage disequilibrium. In this case, alleles at each locus were numerically coded and their inheritance pattern used to group strains that were identical from those that differed at one locus (or share 16 out of 17 markers); isolates that differed at the same genetic locus were grouped together, isolates that differed at a separate locus were connected by a line within each clonal complex. As expected, eBURST clustered the two Type II isolates (2987, 3131) into a single clonal complex with Me49 (CC1). Type X subgroup G isolate 3675, originally identified by its allele at the microsatellite marker ROP1, was included within CC1 because it differed from Type II strains at only SAG4. For the remaining 18 Type X isolates, 15 clustered into 3 clonal complexes (designated CC2-CC4), and 3 unique haplotypes (3671, 3819, 4166) were resolved. The eBURST analysis failed to support a clonal lineage designation for Type X. Further, the increased resolution provided by the 17 markers subdivided the 8 distinct haplotypes defined using 5 loci in Figure 1A into 12 haplotypes (Figure 1D) which was consistent with another study that resolved Type X *T. gondii* genotypes isolated from southern sea otters into 8 genotypes [26]. One isolate each, from subgroups A (n=4; 3168), C (n=4; 3160), D (n=3; 4166) and F (n=3; 3387), was resolved further as unique haplotypes based on the inheritance pattern of alleles at PK1, SAG4 and GRA7 (Supplemental Table 1).

### Type X resembles a recombinant clade of F1 progeny from a natural cross

eBURST is insufficiently resolved to distinguish among isolates that exist either as variants within a clonal lineage (because they possess alleles that differ by only minor mutational drift), or as genetic admixtures that possess genetically distinct alleles derived from different parental types. Previous studies suggested that Type X (or HG12) is a clonal lineage infecting sea otters and wildlife in North America, and that HG12 represents an expanded clone from a sexual cross between a Type II strain and a new genetic lineage that was referred to as “γ”; however this conclusion was based largely on alleles found at just a few loci, including GRA6 and GRA7 [24, 25, 27]. To investigate whether the genetic relationship among the 12 distinct haplotypes is supported by models of genetic drift or recombination, individual maximum likelihood trees were created for each of the 17 nuclear-encoded single copy gene loci, the microsatellite locus, and the two organellar loci (Supplemental Figure 1). To distinguish between alleles undergoing minor mutational drift and those that have evolved independently as distinct genetic outgroups, 1000 bootstrap replicates were run for each tree, with supported nodes above 60% indicated at each marker. For each tree, the 19 isolates identified as Type X were labeled in purple, and isolates belonging to the clonal lineage I (n=1), II (n=3), and III (n=1) were labeled in red, green, and blue respectively (Figure 2).

**Figure 2:**
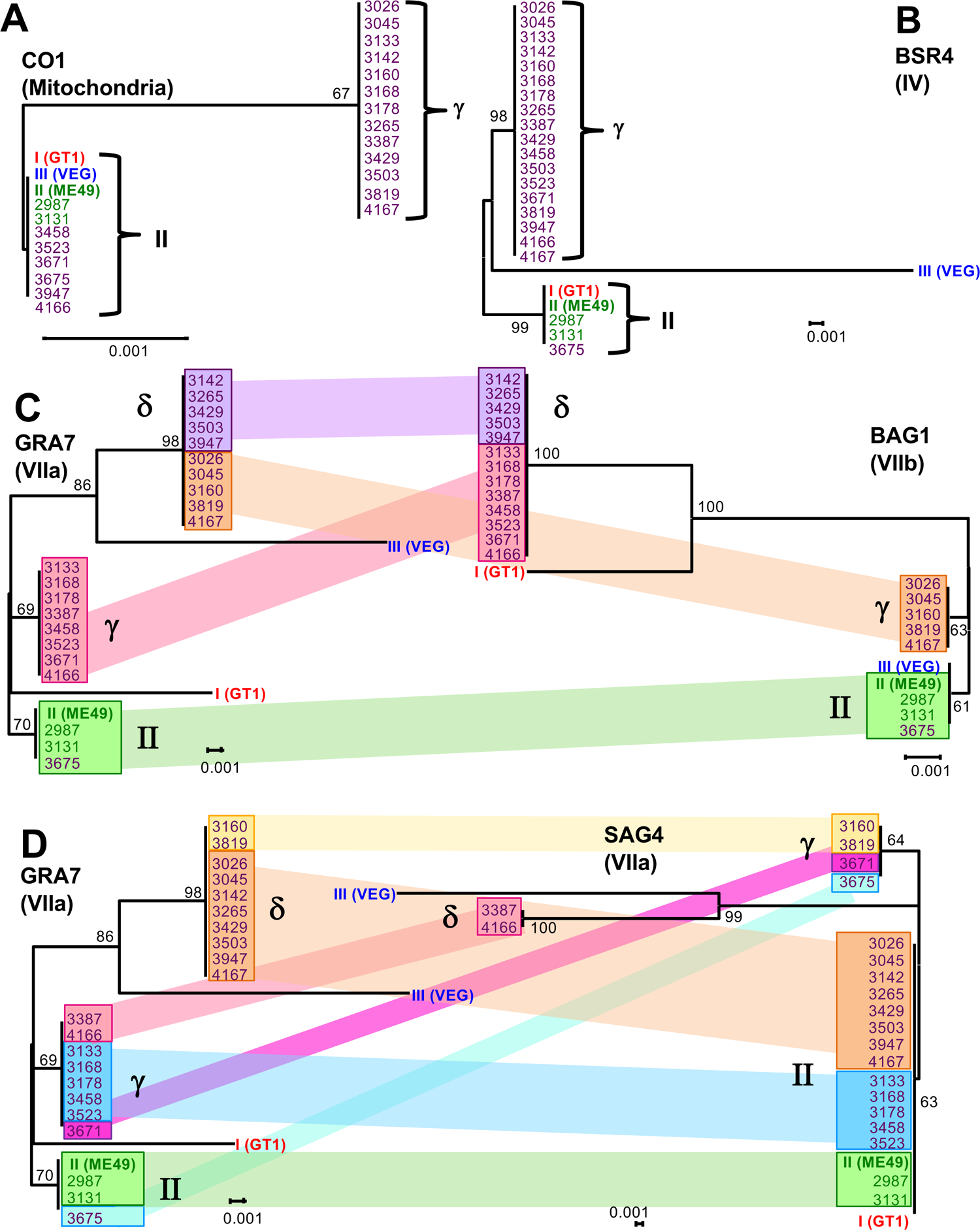
Phylogenetic analyses support designation of Type X *Toxoplasma gondii* isolates recovered from southern sea otters as a recombinant clade of related strains Maximum likelihood trees of sequenced markers are depicted. Allele designations were dependent on a bootstrap support greater than 60%. Isolates are colored based on their genotype from Figure 1A (I: red, II: green, III: blue, X: purple). A) Mitochondrial marker, CO1. B) Nuclear genome marker BSR4 on Chromosome IV. At BSR4, all Type X strains possessed a single allele, referred to as γ that was readily distinguished from Type I, II and III alleles C) Comparison of two unlinked nuclear-encoded markers GRA7 and BAG1. At both markers, Type X strains possessed one of two distinct alleles, referred to as γ or 8. Isolates possessing 8 alleles at both markers were highlighted in purple, those that possessed Type II alleles at both markers were highlighted in green. Isolates that were recombinant and possessed a γ allele at GRA7 but a 8 allele at BAG1 were highlighted in pink, and conversely, a 8 allele at GRA7 and a γ allele at BAG1 were highlighted in orange. D) Comparison of linked genomic markers GRA7 and SAG4. Genetic recombination is highlighted within this chromosome. Isolates that are linked and have Type II lineage alleles at both markers are highlighted in green. Isolates that show recombination between the γ/δ lineage alleles and the Type II lineage alleles are highlighted in pink and orange depending on recombination directionality.

At 5 genetic markers located on chromosomes Ib, II, III, XI, and XII, all 12 distinct Type X haplotypes possessed a canonical Type II allele (green), with no minor mutational drift detected, consistent with Type II being one of the parental genetic backgrounds (Supplemental Table 1). At the L358 marker on chromosome V, however, all Type X isolates except 3675 possessed a Type I or “α” allele (red), indicating the presence of mixed ancestry. Further, at the BSR4 marker on chromosome IV, all Type X isolates except 3675 possessed an entirely novel allele that was readily differentiated from Type I, II and III strains with strong bootstrap support (Figure 2B). At all remaining loci, Type X isolates possessed either a Type II or one of 2 genetically distinct alleles, which could not be readily explained by minor mutational drift. Hence, across all 17 nuclear-encoded markers, each isolate possessed either a Type II allele, or an allele of distinct ancestry that was referred to as belonging to “α” (red), “γ” (purple) or “δ” (orange) lineages.

The maternally inherited organellar genome markers were likewise mixed. At the mitochondrial marker CO1, two alleles were identified, either an allele common to Type I, II and III strains or a novel allele designated “γ” (Figure 2A). Further, 3 distinct alleles were identified at the apicoplast marker APICO: a Type I or “α” allele (red), a Type II allele (green) or a novel “γ” allele (purple), supporting the mixed ancestry designation. Each of the 12 Type X haplotypes were therefore a mosaic of mixed ancestry that possessed some combination of Type II, Type I, “γ” or “δ” alleles that had segregated independently across all loci investigated. The data are consistent with Type X existing as a recombinant clade resembling F1 progeny that was recently derived, without sufficient time to develop minor mutational drift (Supplemental Table 1).

In support of the sexual cross model, incongruity between phylogenies for each haplotype was readily observed between markers located in different portions of the genome. In Figure 2C, 8 isolates shared the same ancestry between the two genetic loci located on separate chromosomes VIIa and VIIb (GRA7 and BAG1 respectively); these variations were shaded either green (Type II alleles at both loci) or purple (γ lineage alleles at both loci). In contrast, 13 isolates had independently segregated chromosomes of mixed ancestry, indicated by orange and pink coloration to depict isolates that were discordant and possessed different ancestral alleles at each locus; the crossing of orange and pink lines highlights this incongruence (Figure 2C). For example, isolate 3671 had a γ lineage allele at GRA7, but a δ lineage allele at BAG1. Recombination within a single chromosome VIIa was also observed at linked markers (GRA7 and SAG4) for 18 strains (Figure 2D). Here, isolate 3387 had a γ lineage allele at GRA7, but a δ lineage allele at SAG4, whereas 3675 had a II lineage allele at GRA7, but a γ lineage allele at SAG4. These results confirm that recombination had occurred and support genetic hybridization as the most plausible explanation for the origin of the 12 Type X haplotypes.

### Type X is an admixture between Type II and a novel genetic background by CGH-array hybridization

A subset of Type X isolates, 2 each from haplotypes A (3142, 3265), C (3026, 3045), F (3503, 3429) and one each from D (3178) and H (3671) were hybridized against a photolithographic microarray that possessed 1517 polymorphic Type I, II and III strain-specific genotyping probes distributed genome-wide [36]. This was done to test whether the MLST genetic markers accurately predicted the presence of large genome-wide haplotype specific blocks, consistent with genetic hybridization. Each Type-specific SNP on the CGH array is represented as a dot colored either red, green, or blue, respectively for a hybridizing strain that possesses a Type I, II or III SNP at the probe position. Grey dots identified probes that failed to hybridize, consistent with strains that do not possess a Type I, II or III specific SNP at the probe position. Hybridization with DNA from a Type I (GT1), Type II (Me49), and Type III (CTG) strain identified hybridization patterns consistent with genotype, indicating that all genotype-specific probes were functioning as expected (Figure 3A). Hybridization with the Type X isolates identified large, contiguous Type II haploblocks, as evidenced by hybridization of all Type II-specific probes present on Chr III, VI, VIIb, and XII, for example. However, in other portions of the genome, the SNP diversity pattern was novel. The hybridization pattern in these regions either shared patch-work similarity with Type I (i.e., at Chr Ib, VI), or was highly divergent, with large contiguous blocks of SNP probes that failed to hybridize (indicated by grey coloration), consistent with introgression of a non-Type I, II or III genetic ancestry, rather than minor mutational drift (i.e., Chr IV, XI, right end of Chr VII, XII, left end of Chr VIII).

**Figure 3:**
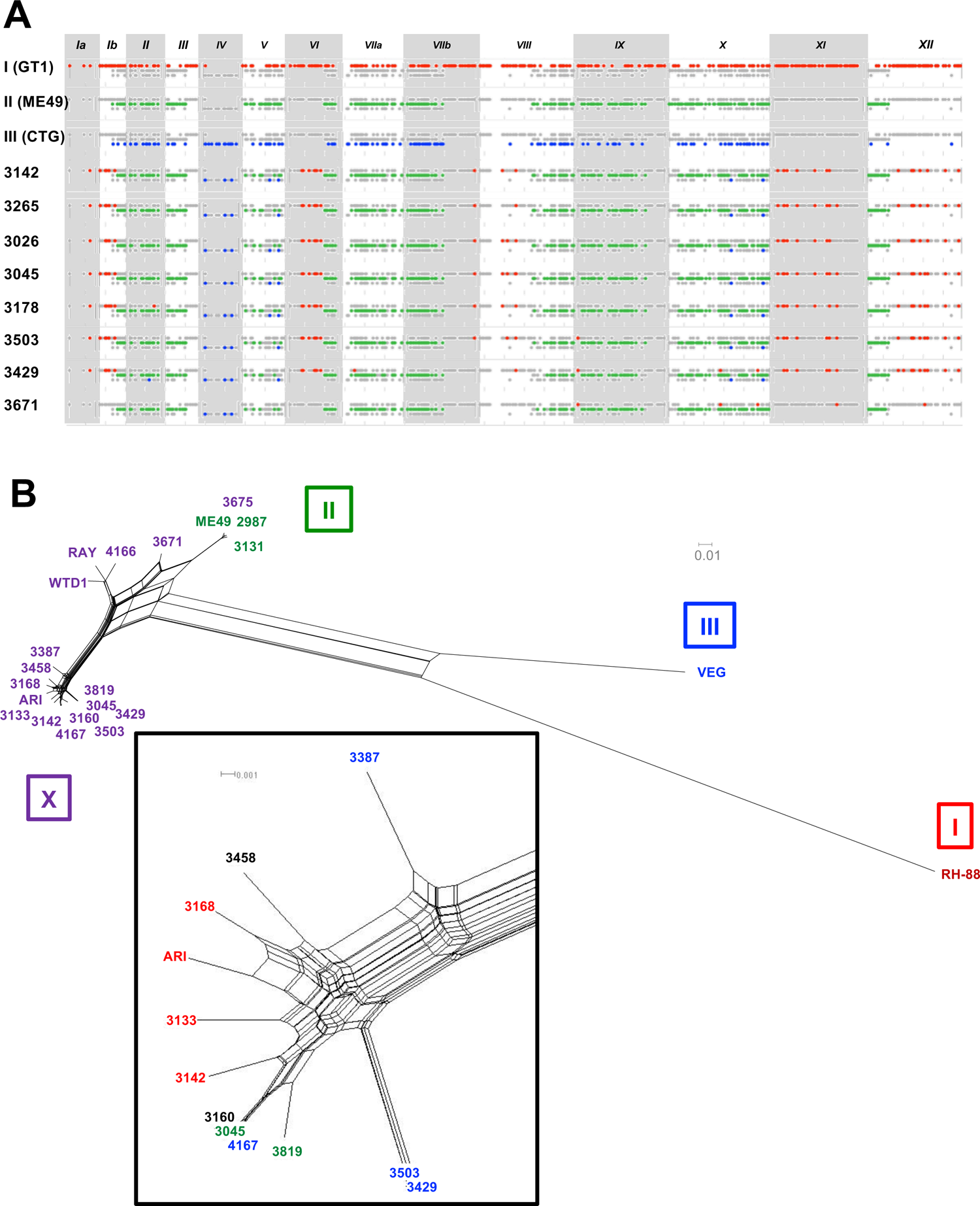
Genome-wide SNP typing of Type X *Toxoplasma gondii* isolates recovered from southern sea otters displays haploblock recombination across the genome A) CGH-array hybridization of Type X strains against Type I, II and III lineage-specific probes identified 8 distinct Type II and non-Type II hybridization patterns across the 8 Type X strains examined. SNPs are represented by a dot in one of three rows indicating where the isolate has hybridized to the microarray with a hybridization characteristic of lineage Type I (red), II (green), or III (blue). Grey indicates no hybridization at this location. Chromosomes are represented as alternating grey and white bars. B) Linkage disequilibrium and network reticulation demonstrates the existence of recombination across the Type X genome. A NeighborNet tree based on whole genome sequence identified 568,592 SNP variant positions across the 19 Type X strains reference mapped to ME49. Type I (red), II (green), III (blue), and X (purple) strains were annotated in the box beside the group based on their previous MLST designation. Two clusters of Type X strains were identified. Strains within the inset were colored based on their murine virulence: virulent (red), intermediate virulent (blue), avirulent (green), not assayed in mice (black).

The pattern of hybridizing SNPs was unique for each of the seven Type X isolates, but it was only minimally different; either 3, 5, or 15 SNP differences were detected in pairwise comparisons within haplotype A (3142, 3265), C (3026, 3045), and F (3503, 3429), respectively (Figure 3A). At this resolution, it was not possible to distinguish between CGH array hybridization efficiency and minor mutational drift. Indeed, the CGH arrays each had approximately the same number of SNP differences (3-15) between haplotypes as they did within each haplotype; this is more consistent with hybridization efficiency as the explanation for hybridization differences. Further, this analysis established that the *T. gondii* isolate 3671 is a novel genetic admixture, because it possessed a distinct Type II hybridization pattern at Chr Ia, the left side of Chr Ib, VI, XI, and right side of Chr XII that differed from all other Type X strains (Figure 3A).

### Type X is a recombinant clade at WGS resolution

Due to the limited number of SNPs and differences in hybridization efficiency using the CGH array approach, WGS was required to infer an accurate genetic history model for the Type X strains. Using genome-wide polymorphism data, 568,592 variable SNP positions were resolved after reference mapping the Type I, II, III, 16 SSO isolates (14 Type X, 2 Type II), and three previously WGS sequenced HG12 isolates (WTD-1, RAY, ARI) against the published Me49 genome. We next performed an unrooted Neighbor-Net analysis (Figure 3B) that resolved the Type X strains into 15 distinct haplotypes at WGS resolution (Supplemental Figure 1) and established that all Type X isolates, similar to Type I and III strains, shared blocks of genome-wide ancestry with Type II, represented by the reticulated pattern of edge blocks that depict recombination events between Type II and the different genetic backgrounds within X, which were distinct from I and III. Our data did not support a clonal lineage designation, as most Type X isolates appeared on separate branches (Figure 3B, Inset). Only clonal complexes 1 and 4 (CC1, Type II; CC4, 3045, 3160, 4167) were supported at WGS resolution, and 3 previously sequenced HG12 strains isolated from 2 people (ARI, RAY) and a deer (WTD-1) formed well-supported clades with Type X isolates 4166 (WTD-1; RAY) and 3168 (ARI), indicating that Type X has a broad host distribution that extends beyond sea otters. Outside of the reticulated network of edge blocks, the branch length for each Type X isolate was significantly less than that observed for Type I and III, synonymous with a more recent origin, without sufficient time to accumulate private SNPs by mutational drift. When murine virulence data for each genotype was mapped onto the Neighbor-Net tree (see inset, Figure 3B), a clear partitioning of the different virulence phenotypes was resolved along branches within the network, consistent with Type X isolates existing as sister progeny, with only minor mutational drift detected at each supported branch.

To identify the size, genome distribution, and total number of Type II admixed haploblocks introgressed into each Type X isolate, pairwise SNP diversity plots were generated for all Type X isolates, as well as reference Type I and III strains that have been previously shown to have recombined with Type II. The Type I GT1 strain possessed an Me49 Chr Ia and IV, and an Me49 admixture block at the left side of Chr VIIa, and right end of Chr XI (Figure 4). As expected, the Type III VEG strain possessed many more Me49 admixture blocks distributed genome-wide than GT1, comprising ∼40% of its genome. Only Type X haplotype H (3671) possessed haploblocks that were highly similar in sequence to Me49, on Chr Ia, II, III, VI, VIII, IX, XI, and XII (Figure 4). In other regions, however, 3671 shared regions clearly Me49-like, but that appeared to have diverged somewhat by mutational drift (average 3-5 SNPs per 10kb block) on Chr II, or introgressed into Chr IV, V, VIIa, VIIb, VIII, IX, and XII. Intermixed within the Type II regions of the 3671 genome were a limited number of large haploblocks containing divergent SNP density synonymous with hybridization by a strain possessing distinct γ or δ genetic ancestry (average of 50-100 SNPs per 10kb block). When the pairwise SNP density analysis was expanded to different Type X haplotypes, isolates within each haplotype possessed highly similar patchwork mosaics of Type II-like or divergent (Type X) haploblocks that were specific to each haplotype. For example, haplotype A isolates 3142 and 3168 SNP density plots were highly similar to each other, and to haplotype C isolate 3045, but were readily distinguishable from isolates 4166 and 3503; strains within haplotypes D and F, respectively (Figure 4). Specifically, isolate 3142 had a Type X haploblock inheritance pattern at the right end of Chr V, and in the middle of Chr Ib, VIIa, VIII, IX and XII, whereas 4166, which was indistinguishable from a previously sequenced HG12 strain (RAY) recovered from a human patient, possessed either Type II or 3671 haploblocks in these regions (Figure 4). Although the haplotype F isolate 3503 was highly similar in genomic organization to 3142, it possessed a Type II haploblock at the right end of Chr V that readily distinguished it from haplotype A. The pairwise SNP analysis established that most Type X haplotypes possess a genomic architecture that is strikingly similar, but different at a limited number of admixture blocks. Coupled with the low allelic diversity, the data support a genetic history model whereby Type X resembles a sexual clade of recently-derived natural recombinants from a relatively limited number of crosses.

**Figure 4:**
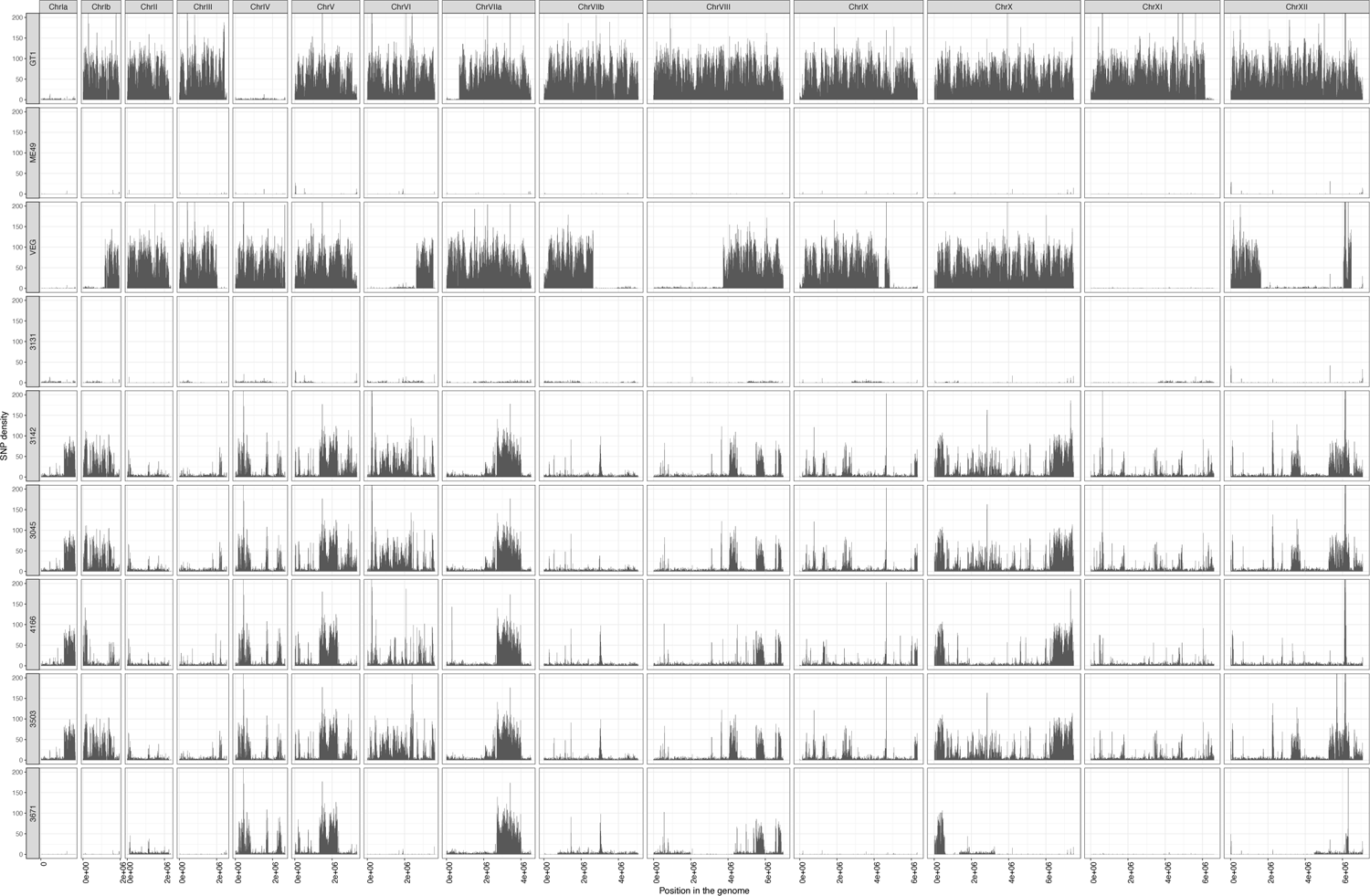
SNP density plots for *Toxoplasma gondii* isolates recovered from southern sea otters identify distinct and different recombination crossover points between the Type II and γ/δ lineage among examined Type X strains WGS reads for all strains were mapped to the ME49 reference genome. Each row represents the SNPs of one strain mapped in a sliding 10 kb window across the genome, separated into chromosomes columns. Vertical bars across the row represent the number of SNPs, the height of the bar (from y-axis 0-200 bp) represents the amount of divergence from the ME49 genome. Type X strains were grouped based on their MLST designation.

### PopNet analysis identifies only limited admixture blocks among Type X isolates

While the Neighbor-Net analysis established that Type X exists as a recombinant clade of strains, it failed to predict the precise number of genetic ancestries, or how they had admixed positionally across the chromosomes. PopNet was used to paint chromosomes according to their local inheritance patterns [37]. Included in the analysis were all sequenced Type X and II strains, as well as reference Type I and III strains known to have admixed with Type II. PopNet identified 4 statistically supported ancestries and showed that each genome was a mosaic of these distinct ancestries, further supporting the admixture model.

Within the circle depicting the Type X clade, 5 distinct subgroupings were identified based on the number and position of shared ancestral blocks; these were grouped together based on line thickness (Figure 5A). These same groupings were supported both by Pairwise SNP plots and Neighbor-Net analysis, but the PopNet analysis showed which of the 5 ancestries had introgressed positionally across the mosaic genomes. Clear recombination blocks were readily resolved, and the genome architecture was remarkably similar between the groupings. A custom script designed to identify only major crossover points between the two parents (Type II and the mosaic ancestry of the γ/δ parent) identified either a limited number of single and double recombination events, or the inheritance of whole chromosomes of either Type II (Chr II) or γ/δ (Chr Ia, IV, XI) parental ancestry, consistent with this group of strains representing related sister progeny (Figure 5B). Because murine virulence plotted on the Neighbor-Net tree identified clear pathogenicity differences in mice that clustered based on parasite genotype, and that the Type X strains resembled a natural clade of recombinant progeny (akin to F_1_) that possessed only a restricted number of crossover points, a population-based QTL was performed to see if genes could be mapped that contributed to murine virulence within the set of 18 natural isolates for which virulence data existed (Figure 5C),

**Figure 5:**
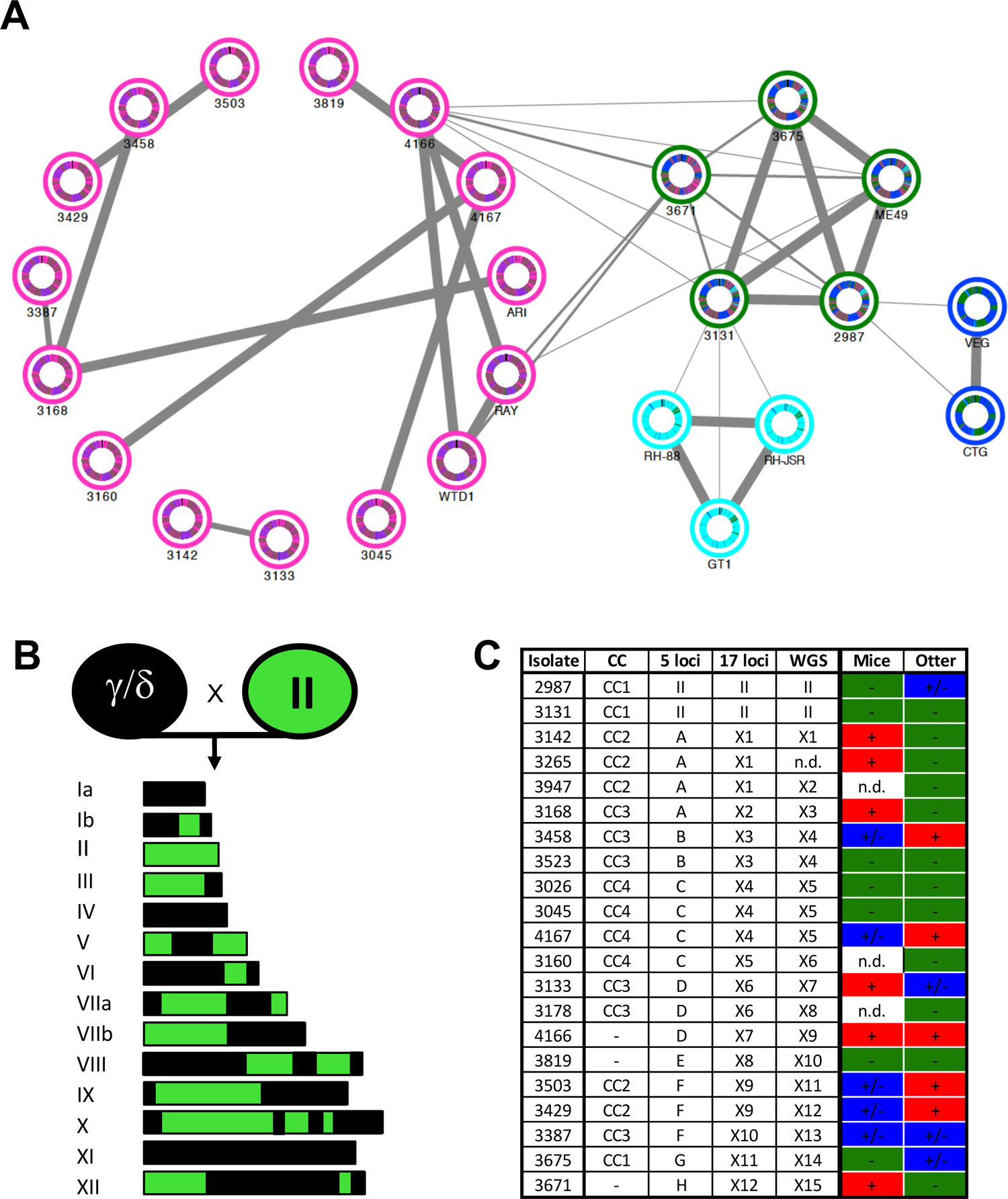
Whole genome sequencing establishes *Toxoplasma gondii* Type X isolates recovered from southern sea otters as a recombinant clade of related strains A) PopNet analysis for all strains sequenced at WGS resolution. The genome of each strain is represented by a circle of concatenated chromosomes starting at the top with Chromosome Ia and rotating clockwise to Chromosome XII. All strains were clustered into four distinct color groups based on their degree of shared ancestry (Type II-green, Type X-pink, Type I-cyan, Type III-blue) and each isolate has an inner circle that is arrayed in color-group haploblocks based on shared ancestry. Line thickness between strains indicates their interrelatedness, boldest lines indicate 90-100% of the genome is in linkage disequilibrium, as shown for strains 4167, 3819, 3045 and 3160. B) Genetic model showing cross-over recombination points for isolate 3142. Type II ancestry is shown in green whereas the mosaic of γ/δ ancestry is shown in black. C) 21 WGS isolates are shown with their group designation for the 5 loci MLST (Figure 1A), eBURST clonal complex identity (Figure 1D), 17 loci MLST (Supplemental Table 1), and murine virulence phenotype (Figure 1C).

### Natural Population-based QTL identifies ROP33 (WNG-3) as a new murine Vir locus

All Type X isolates recovered from sea otters resembled offspring from one (or a few crosses) between a Type II strain, which is avirulent in mice, and unknown parent(s) that are a mosaic of two distinct ancestries. Because these natural isolates possessed a range of virulence phenotypes in laboratory-exposed mice (Figure 1C), a genome scan was performed to determine the log-likelihood for association of discrete genome haploblocks with the acute virulence phenotype. Four quantitative trait loci (QTL) peaks were identified with logarithm of odds (LOD) scores 4.0 or greater on chromosomes V, VIIa, VIII, and X (Figure 6A). The average size of the genomic regions spanned by the QTLs were in the range of 100-200kb, except for one on chromosome V, which was >700 kb (Figure 6B). To identify candidate genes within the four peaks, the following inclusion criteria were assessed: presence of a signal peptide and/or transmembrane domain, gene expression and polymorphism differences, as well as every gene’s CRISPR genome-wide mutagenesis score for essentiality [38] (Supplemental Table 2). *ROP33* on chromosome VIIa stood out as the best candidate gene to target for reverse genetics as it was abundantly expressed during acute infection, it was predicted to be a functional serine-threonine protein kinase, it was highly polymorphic, and is part of a family of divergent WNG (with-no-Gly-loop) kinases that regulate parasite tubular membrane biogenesis [33]. Further, one allele of *ROP33* was strongly correlated with acute virulence in mice (p=0.00031; fishers two-sided exact test) (Figure 6C).

**Figure 6:**
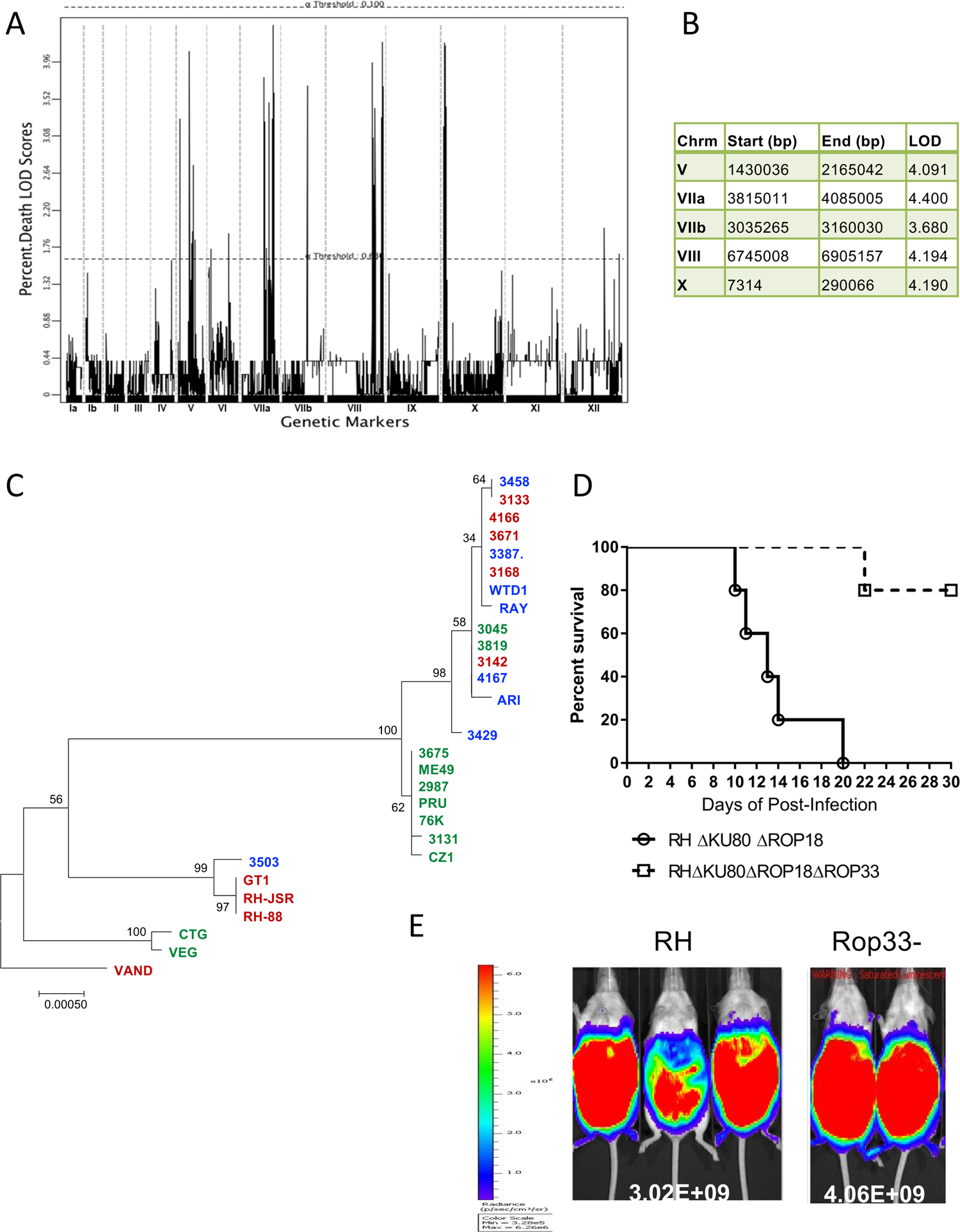
Population level QTL analysis identifies multiple chromosomal locations associated with acute murine virulence A) WGS data from Type X strains was down-selected to one SNP every 5 kb and analyzed using standard QTL methods with 1000 bootstrap support. LOD scores are shown across the chromosomes for these strains based on genetic association with acute murine virulence as shown in Supplemental Table 2. B) Significantly associated peaks from the QTL were identified based on clusters of SNP peaks. The LOD scores for the tallest portion of the associated genomic region are listed. C) Distance-based Neighbor-joining ROP33 tree. Alignment of *ROP33* DNA sequences were used to construct a phylogenetic tree representing the majority of ROP33 alleles encoded within the species *Toxoplasma gondii.* Mouse virulence for each isolate is depicted by a color-code, per Figure 1C. **Red**, mouse virulent, **Blue**, intermediate mouse virulent (some CD-1 mice survived acute infection); **Green**, mouse avirulent (all CD-1 mice survived acute infection). Inoculum size was 50 tachyzoites injected intraperitoneally in CD-1 outbred mice. D) Type I RH parasites deficient in ROP18 and ROP33 are less virulent than Type I RH parasites deficient in ROP18 alone, indicating that ROP33 is a virulence locus in *Toxoplasma.* CD-1 mice were infected intraperitoneally with 500 tachyzoites of the RH*Δrop18* or RH*Δrop18Δrop33* DKO, only the results for seropositive mice are shown. Data are combined from two independent experiments, each with five mice per group. E) Bioluminescent detection of parasite burden *in vivo* at 9 days post-infection of CD-1 mice with 50 tachyzoites injected intraperitoneally. A representative image with photon output in photons/second/cm2/surface radiance (*sr*) is shown.

To determine whether *ROP33* was a novel murine virulence gene, the *ROP33* locus was disrupted by targeted deletion using CRISPR-Cas9 facilitated double crossover homologous recombination in the RHΔ*ku80*Δ*rop18*Δ*hxgprt* strain [39]. This strain is virulent in mice (LD_100_=500 tachyzoites) and was engineered to accept targeted replacement of the *rop33* gene using an HXGPRT gene flanked by 30bp of homology just outside of the *rop33* promoter and 3’UTR region. Following selection in mycophenolic acid (MPA) and xanthine (to select for the *HXGPRT* gene), the population was screened for disruption of the *rop33* gene by PCR. Mice were next infected with either the parent RHΔ*ku80*Δ*rop18*Δ*hxgprt* strain or the Rop33^-^ mutant to assess virulence. Groups of 5 outbred CD1 mice were injected intraperitoneally with 500 tachyzoites and the results shown are for one of two replicates. All mice infected with the parent RH line succumbed to infection within 20 days (Figure 6D). In contrast, targeted deletion of the *rop33* gene was protective, with most (9/10) mice surviving acute infection. Our results suggest that ROP33 is an acute virulence gene for *T. gondii*. Importantly, the lack of virulence for parasite strains with deletion of the *rop33* gene was not the result of a failure of the mutant parasite to proliferate *in vivo*, as no difference in parasite load was detected through acute infection by bioluminescence imaging (Figure 6E). All surviving mice were seropositive for *T. gondii*, indicating that they had been productively infected. These data suggest that ROP33 is a new virulence factor for murine infection and that this mouse-virulence gene was identified from wild-derived *T. gondii* isolates that had undergone a population genetic cross in a natural setting.

## Discussion

We investigated the emergence of a suspected clonal lineage of *T. gondii* (Type X; also referred to as HG12) from isolates recovered longitudinally from a single marine host (southern sea otters) in California over seven years (1998-2004). Our goal was to determine the genetic basis for the emergence of this unique *T. gondii* lineage in a federally listed, threatened species. Although Type X had previously been identified as the 4^th^ clonal lineage in North American wildlife, our work established that Type X is not a clonal lineage, but rather a recombinant clade of strains that resemble F_1_ progeny from at least one natural cross. DNA sequence analysis using diverse linked and unlinked markers across the nuclear and organellar genomes of Type X isolates identified Type X to be a composite of Type II, and a mosaic of two distinct ancestries, referred to as γ and 8, that had recombined to produce a highly invasive clade of strains causing both subclinical infection and mortality in a threatened marine mammal.

Our data suggest that Type X was derived from one or a limited number of natural crosses between two highly related parental strains. One sea otter genotype (Type X haplotype A) was widely distributed and was found to have expanded to cause most subclinical infections in sampled sea otters, but was highly virulent to laboratory-exposed mice. Our isolates from a naturally-infected wild animal population support a model whereby *T. gondii*’s sexual life cycle is facilitating the evolution and expansion of cryptically virulent strains that can cause significant morbidity and mortality in a species-specific manner. For generalist parasites like *T. gondii* that have broad host ranges, natural selection among intermediate hosts appears to maximize transmission and may establish how virulent strains can be maintained cryptically in nature. The expansion of a mouse virulent clone into a new ecological niche, such as nearshore marine mammals of the Eastern Pacific coastline, demonstrates how *T. gondii* is leveraging its intermediate host range to selectively partition parasite genetic diversity and epidemic disease potential.

In prior studies, unknown strains having a distinct ancestry, referred to as α and β, were found to have crossed with Type II to create the Type I and Type III clonal strains, respectively [11]. The frequency and pattern of recombined blocks inherited within the Type X strains is parsimonious with previously described models for the genetic history of Types I and III in which an unidentified ancestor (respectively α and β) sexually recombined with an ancestral Type II to create new lineages (Figure 5B) [11, 40]. Prior studies have suggested that Type X is a product of genetic hybridization between a Type II lineage and an unknown γ lineage [25, 27, 28, 32, 41]. In the current study, we have further refined this perspective and concluded that Type X is a mosaic of a cross between Type II strains and two distinct γ and δ ancestries. The γ/δ lineage has likely recombined at least once with Type II to create the Type X recombinant clade of strains. Currently, no single isolate has been found which could be defined as either γ or 8, similar to the α and β lines thought to have admixed with Type II to produce the clonal lines I and III [11, 40].

Previous work on Type X circulating in wildlife samples using a limited set of genotyping markers, including sea otters, identified Type X as the 4^th^ clonal lineage in North America [27, 28]. However, this designation was based on the discovery of genetically distinct alleles at only a few loci, including GRA6, B1 and SAG1, which was the original basis for distinguishing the Type X lineage as distinct from Type II, and that it had undergone a hybridization event with a novel genotype [25]. It is now well established that Type X strains commonly infect wildlife across North America [14, 26–30, 32, 35]. This study examined *T. gondii* strains isolated longitudinally across a 7 year period and a 680 km expanse along the California coast from stranded southern sea otters. Our sample set was comprised of otters with *T. gondii* infections that ranged from subclinical to fatal.

Because previous work had classified Type X as clonal, detection of multiple disease states among infected sea otters was unexpected [27, 28, 32, 35]. To ascertain whether parasite genotype was influencing the observed disease spectrum, genotyping studies were carried out using an expanded set of markers to more definitively characterize the population genetic structure of Type X isolates. All genotyping markers used in this study were sequenced to increase resolution rather than evaluated strictly for their PCR-RFLP genotype, as was standard for many prior studies. While the PCR-DNA-Seq markers used herein only surveyed a small part of the genome, they established that strains genotyped as Type X via GRA6 and SAG1 alleles were in fact comprised of at least 12 distinct haplotypes that could not be classified as minor variants based on relatively few SNP differences, as concluded previously [26]. Furthermore, at any given locus, only a Type II allele or one or another of two distinct alleles was identified, indicating limited allelic diversity and a total lack of private SNPs. Hence, each of the 12 haplotypes possessed one of just three allelic types at any one locus that had independently segregated across all examined markers. This result is parsimonious with a recombinant clade of strains that reflect genetic hybrids from a limited, but distinct set of ancestries.

Our results were further confirmed both by CGH array analyses and whole genome sequencing. The previous misclassification as a clonal population appears to be the result of the low resolution genotyping analyses performed, and the placement of the markers in predominantly Type II regions of the genome [14, 28, 42]. For this reason, future studies should use WGS to discriminate between haplogroups within a population and determine the true genetic ancestry of each haplogroup. *Toxoplasma gondii* population genetics would also greatly benefit from examination of an increased number of isolates at whole genome sequencing resolution. Hence, WGS should be expanded to include a greater diversity of strains, and multiple isolates from within each canonical clonal lineage.

The Type I and III clonal lineages are thought to be derived from a limited number of sexual crosses, and all strains within these clonotypes share large haploblocks of Type II-like sequence [11, 41–43]. This finding also holds true for the Type X lineage. At all MLST loci surveyed, all Type X isolates possessed at least one marker that was Type II, and the introgression of large haploblocks of Type II ancestry across each genome was confirmed in the 19 strains that were resolved at WGS resolution. In regions that did not clade with Type II, two different allelic types were identified. We concluded that these represented distinct ancestries that we referred to as γ and 8. This designation was supported by the inherent reticulation of the clade, seen in the NeighborNet tree (Figure 2B). Predominantly short branch lengths radiating from a reticulated network with multiple strains on a single branch were resolved, rather than a star-like phylogeny with multiple alleles present at each major branch (the result of private SNPs that accumulated through genetic drift). In fact, the lack of genetic drift between these loci supported genetic hybridization as the most parsimonious explanation for the relationship between the Type X haplotypes.

Sexual replication with genetic recombination is known to occur in *T. gondii* strains in South America at high frequency [40, 42, 44]. Recent studies on the population genetic structure of *T. gondii* have also established that genetic hybridization, at whole genome resolution, is extant for most sequenced strains that represent the breadth of genetic diversity within *T. gondii* [15, 37]. Evidence of sexual recombination across the population, and evidence of sexual replication within the Type X clade shown here, may indicate that sexual recombination is occurring more frequently than previously suspected among *T. gondii* strains circulating within North America.

No barriers appear to exist, or have been described, to limit sexual recombination of *T. gondii* from occurring in a laboratory setting, although there is incomplete understanding of the genetic factors regulating sexual replication in felid hosts [7, 43, 45–47]. It is not clear why sexual recombination is reported less commonly in North American wildlife, when compared to South America, although it is possible that self-mating within dominant populations in North America is masking the detection of genetic outcrossing [12, 41, 48]. Among closely related isolates, unisexual mating (defined as intra-lineage crosses between similar strains that possess distinct genotypes) cannot be resolved using current genotyping methods that rely on low-resolution analyses, and thus cannot readily distinguish asexual expansion from unisexual mating [12].

*Toxoplasma gondii* has been shown to utilize its sexual cycle to expand its biological potential and alter its pathogenesis [7, 45, 47]. Sexual recombination, by the ability to reshuffle parental alleles into new combinations, can produce progeny with diverse biological potential [7]. Our work identified a single *T. gondii* genotype (Type X haplotype A) that was presumably derived from a recent genetic cross and naturally selected in southern sea otters. Although most haplotype A-infected sea otters had mild infections, this genotype was highly pathogenic to outbred laboratory mice, with all mice dying in 10-15 days after administering as few as 50 tachyzoites. In contrast, three Type X haplotype F isolates were highly pathogenic to sea otters but significantly less so to outbred laboratory mice (Supplemental Figure 1). Our data support a virulence shift model for the expansion and propagation of *T. gondii* in nature, whereby the parasite utilizes its sexual cycle to generate a diverse spectrum of new genotypes that possess altered biological potential that can be naturally selected across *T. gondii*’s vast host range to optimize parasite transmission, and colonize new niches. Hence, natural selection among intermediate hosts allows *T. gondii* to maintain highly pathogenic strains cryptically; these strains can cause serious disease in some animal hosts, but not others.

This type of selection has been observed previously in laboratory mice that express variable IRG (Immunity Related GTPase) gene arrays. IRGs are host proteins primarily responsible for combatting *T. gondii* lysis of murine cells. The *T. gondii* genome encodes a suite of highly polymorphic rhoptry kinase genes (ROPKs) that function to inactivate host IRGs [49–51]. While laboratory mice are highly clonal, wild mice and their subsequent IRGs are more diverse, as are the ROPKs expressed by *T. gondii* strains that are capable of infecting wild mice [17, 52]. As a result, different *T. gondii* genotypes are naturally selected in wild mice based on the combination of host IRGs versus ROPK alleles expressed. Hence, wild CIM mice do not succumb to *T. gondii* Type I infections, whereas laboratory mice do.

Similarly, TLR 11/12 is a primary rodent innate immune response sensor (otherwise referred to as PAMPs) that detects *T. gondii* infection. Not all intermediate hosts share a functional combination of TLR11/12; *T. gondii* strains that are selected for mouse infection must bypass TLR11/12 recognition, while this is not a barrier to infection of human cells [52, 53]. As a result, intermediate hosts are capable of naturally selecting *T. gondii* strain genotypes that establish chronic, transmissible infections in a host-specific and parasite strain-dependent manner. To optimize its biological potential, the parasite must find a balance between virulence, fitness, and infectious transmissibility across a wide host range. It is this capacity to maintain cryptically virulent strains that may contribute to *T. gondii’s* global distribution and success in nature; it has the ability to cause disease outbreaks, expand its host range into new ecological niches, or alter its pathogenicity in both a parasite strain-specific and host-specific manner to maximize transmission.

For other protozoan parasites, host partitioning is usually synonymous with speciation. For example, *Plasmodium* spp. often partition by the mosquito host species they co-evolved with to maximize parasite transmission [54–58]. Likewise, *Sarcocystis* spp. largely maintain separate parasite species for each intermediate-definitive host species combination to maintain its life cycle [12, 59]. In contrast, *T. gondii* forms transmissible cysts in virtually all warm-blooded vertebrates, and these cysts are infectious for its definitive felid host as well as diverse intermediate hosts. The difference is that across the genetic diversity of *T. gondii*, specific parasite genotypes are being selectively expanded within different intermediate hosts, based primarily on parasite genotype and the suite of polymorphic effector proteins encoded by each parasite strain. This allows *T. gondii* to maintain cryptically virulent strains across a broad intermediate host range, with intermediate hosts playing a central role in the natural selection, expansion, and maintenance of virulent strains across the host range of this generalist parasite. In effect, infected hosts can act as reservoirs for the pathogenic or epidemic potential of the species.

Our study showed that one sub-type of *T. gondii* which had expanded in most sea otters (where it caused primarily subclinical, chronic infections) was uniformly lethal to outbred laboratory mice. This is similar to another sub-type of *T. gondii* that express predominantly Type I alleles and chronically infect wild birds which is highly pathogenic in exposed laboratory mice [60]. Additionally, while Type II isolates commonly infect domestic livestock in North America, Type X is more common in sylvatic hosts, prompting the hypothesis that separate cycles exist within the *T. gondii* population that overlap solely within the feline definitive host [10, 29, 61].

Finally, only a limited number of crossovers were detected among strains that were WGS sequenced within the recombinant clade of Type X haplotypes. Because the pedigree of one of the parents was known, we concluded that the isolates resembled F_1_ progeny from a natural cross, which prompted us to perform a population-based QTL analysis on an unmanipulated, natural population of genetic hybrids. Taking advantage of a genome-wide, high resolution SNP map we had generated for all sequenced haplotypes, our analysis identified multiple punctate QTL peaks containing a limited number of candidate loci associated with differences in mouse virulence for reverse genetic follow-up. The serine-threonine protein kinase ROP33 stood out as the best candidate to influence pathogenicity; it was previously identified as an active protein kinase that is both abundantly expressed and highly polymorphic. Further, ROP33 is a divergent WNG kinase (WNG-3) that is related to ROP35 (WNG-1); a critical regulator of tubular membrane biogenesis, the formation of the parasite’s intravesicular network (or IVN), and the phosphorylation of dense granule proteins associated with IVN biogenesis [33]. In fact, dense granule proteins GRA2 and GRA6 are required for IVN biogenesis, and IVN-deficient parasites grow normally *in vitro*, but are attenuated in mouse virulence assays [62, 63]. Related ROP proteins, including ROP 5, 16, 17, and 18 are known to hijack host immune signaling pathways to alter *T. gondii* virulence during rodent infection.

However, all Type X isolates investigated in our study, whether mouse virulent or avirulent, only expressed avirulent allele combinations of the known ROP 5/16/18 virulence factors. We chose to knock-out the gene in a Ku80 deficient strain of *T. gondii* that was both mouse virulent, and deficient in ROP18 because no change in mouse virulence was reported when ROP33 was deleted in a wild-type RH strain background [64]. In addition, previous genetic mapping studies failed to identify the QTL for this locus, even though it is polymorphic between Type I, II and III strains for which genetic crosses have been performed. It is likely that the ROP5 and ROP18 virulence factors were dominant, and ROP33 is analogous to ROP17; another virulence factor that was only identified after the virulence enhancing capacity of ROP18 was removed [39].

Our results show that parasites deficient in ROP33 in the highly permissive RHΔku80Δ*rop18* parent were avirulent, whereas an infectious dose of 500 tachyzoites of the RHΔku80Δ*rop18* parent genotype was uniformly virulent in mice. Specifically how ROP33 contributes to murine virulence, what dense granule proteins it phosphorylates, the integrity of the IVN and what host immune signaling pathways are modified are being actively investigated. Future work will dissect the host or parasite factors that ROP33 targets to reduce or influence parasite transmission and pathogenicity. Finally, our study establishes that it is possible to use *T. gondii* isolates derived from a naturally-occurring population genetic cross to identify genetic loci associated with a specific quantitative phenotype.

## Materials and Methods

### Parasite culture

Fifty-three previously published *T. gondii* strains isolated from brain and/or other tissue samples from stranded southern sea otters were provided by Dr. Patricia Conrad [25, 26]. Aseptically-collected tissue for parasite isolation was obtained during gross necropsy of sea otters at the Marine Wildlife Veterinary Care and Research Center in Santa Cruz, CA. Parasite isolation in tissue culture was performed at the University of California, Davis School of Veterinary Medicine, as described [25]. The primary and any contributing causes of death (COD), sequelae, and incidental findings were determined for each sea otter based on gross necropsy and histopathology, as described [25, 26, 31, 65]. The coastal location for each stranded sea otter, as well as the associated *T. gondii* isolate, was noted to the nearest 0.5 km along the central California coast as ATOS (“as the otter swims”) numbers. Isolates were named sequentially based on the sea otter number and stranding year. Due to multiple encounters with the same otter over its lifetime, sea otter case number 3142 was later corrected to 2922, but both numbers represent the same animal; this is indicated in Supplementary Figure 1. Parasites were maintained in vitro in human foreskin fibroblast (HFF) monolayers as described [66].

### DNA extraction and genetic typing markers

Cultivated parasites were syringe lysed using a 27-gauge needle and filtered through a 3.0 micron polycarbonate filter to remove cellular debris. DNA was extracted from cell pellets using the Qiagen DNeasy Blood and Tissue kit (Qiagen). All 53 isolates were typed using 5 previously described PCR-RFLPs, and one DNA sequencing marker [7, 25, 67–69]. BSR4, BAG1, and ROP1 were PCR amplified and typed based on known RFLP identity. GRA6 and SAG3 were typed according to previously identified representative SNPs based on Sanger sequence data by the National Institute of Allergy and Infectious Disease’s Rocky Mountain Laboratory (NIAID RML).

Alleles observed via sequencing were designated as canonical I, II, or III. When sequences differed from canonical alleles, previously described methods of characterization were used [29]. Briefly, when genetic variation differed from references by two mutations or more, the allele was classified as a novel allele, whereas only one mutation was labeled as a drifted allele from the closest reference sequence.

Twenty-one strains representing the δ unique genotypes identified by the 5 markers applied herein were then selected for additional characterization. These 21 strains were PCR amplified, sequenced, and genotyped using 20 markers: 18 at both linked and unlinked genomic loci, encompassing 13 of the 14 chromosomes, and 2 loci on organellar genomes (apicoplast and mitochondria), representing 15,430 bp with 335 SNPs (∼0.024% of the genome). Sequences were examined and nucleotides were verified using SeqMan Pro alignment software (Lasergene). Sequences for reference strains (ME49, GT1, and VEG) were downloaded from ToxoDB [70].

### Generation of the *ROP33* knockout strain in *RHΔku80Δhxgprt* parasites

Deletion of the *ROP33* gene was generated using the Type I parasite RH that was rendered deficient in the expression of *KU80* and the drug selectable marker *HXGPRT*. Briefly, the CRISPR/Cas9 targeting plasmid pSAG1:CAS9,U6:sgUPRT (received from Prof. L. David Sibley, Department of Molecular Microbiology, Washington University School of Medicine) was modified with a guide RNA (GTCGGACGCGAAACTCGCTT) to target the 5’ region of the *ROP33* gene (sgROP33) using Q5 mutagenesis (New England Biolabs, MA). Then a CRISPR/Cas9 replacement construct was created using Gibson assembly (New England Biolabs) to stitch together a 1kb flanking region of the 5’ UTR region of ROP33 (TGGT1_chrVIIa, 3921773 to 3922773 bp) with the selectable marker HXGPRT and a 1kb flanking region of the 3’UTR region of ROP33 (TGGT1_chrVIIa, 3926189 to 3927189 bp) to replace the *ROP33* gene using the selectable drug cassette. To delete the *ROP33* gene, a total of 50 μg of the sgROP33 and replacement construct (5:1 ratio) were co-transfected into a ROP18 deficient strain of RH (RH *Δku80ΔhxgprtΔrop18*), which was cultured in human foreskin fibroblasts (HFF) cells and maintained in Dulbecco’s modified Eagle’s medium (DMEM) supplemented with 10% fetal bovine serum, 2 mM glutamine and 25 μg / ml gentamicin. After transfection, parasites were selected for stable integration of the targeting construct using mycophenolic acid (12.5 mg / mL in MeOH; MPA) / xanthine (20 mg / mL in 1M KOH; XAN) as described [71]. After 15 days, the resistant population was cloned by limiting dilution and single clones were screened by PCR for targeted deletion of the *ROP33* gene.

### Mouse virulence assay

To determine the virulence of the sea otter *T. gondii* isolates in a mouse model, groups of five or more, 6-8 week-old, female CD1 outbred mice were intraperitoneally injected with 50 *T. gondii* tachyzoites resuspended in 500 μl of PBS that had been expanded in HFF cells. Mice were weighed daily to measure infection-induced cachexia, and mouse survival was assayed over 42 days [7]. At 14 days, mice were bled, and serum was extracted to test for seroconversion via indirect fluorescent antibody test (IFAT) against ME49 tachyzoites [72]. Strain virulence was classified as follows: avirulent strains killed no seropositive mice within 42 days of infection, intermediate virulent strains killed some but not all seropositive mice within 42 days post-infection, and virulent strains killed all seropositive mice. To assess the contribution of ROP33 to mouse virulence, groups of five, 6-8 week old female outbred CD1 mice were injected intraperitoneally with 500 tachyzoites of either the parent RHΔ*ku80*Δ*rop18*Δ*hxgprt* strain, or the *Rop33^-^* mutant; infectivity and virulence were assessed as above.

### Phylogenetic Tree and DNA Marker Analysis

The default settings of Clustal X were used to align DNA sequences for all markers individually and as a concatenated set [73]. MSF files of aligned marker sequences were imported into Molecular Evolutionary Genetic Analysis (MEGA) Version 7 to create a maximum likelihood tree using Tamura-Nei model distance analysis with uniform rates of substitution across all sites, and 1000 bootstrap support for all branch points [74]. Consensus trees were rooted on ME49, as Type II is an inherent parent in all strains analyzed. All scales were set at 0.001 nucleotide difference unit. Distinct parental lineages with bootstrap support over 60% were indicated on each tree to distinguish II alleles from novel γ and δ alleles. These novel alleles were indicated in purple and orange respectively for the 17 MLST loci (Supplementary Table 2). The Nexus file of the concatenated marker alignment was imported into SplitsTree 4 Version 4.13.1 where NetworkNet analysis was run using default settings [75].

### eBURST Clonal Complex Analysis

Alleles determined from the 17 nuclear encoded markers (excluding ROP1 due to its classification as a microsatellite marker) were given numerical designations to create a multilocus sequence-typing scheme for the eBURST program [76]. Default settings were used to evaluate strain-specific alleles. Individual strains were displayed as dots with unique colors: Type I, II, III, and X strains were represented by red, green, blue, and purple dots respectively, and dot size reflected the number of strains assessed within each genotype. Strains sharing 16 of the 17 nuclear encoded markers were designated as clonal complexes, represented by interconnecting lines in the diagram.

### Genomic hybridization to Affymetrix arrays

Genomic DNA was sheared and biotin labeled before hybridizing to a custom *T. gondii* Affymetrix microarray as previously published [36]. High-fidelity SNPs were characterized via a custom R script to identify SNPs belonging to each of the three reference strains (I, II, and III). Three reference strains (GT1, ME49, and CTG) were shown to demonstrate ideal hybridization within canonical lineages. Sea otter Type X DNA isolate hybridizations were shown below the reference strains.

### Whole-Genome Sequence Analysis of Type X Recombination

DNA was isolated from 16 southern sea otter *T. gondii* isolates with optimal tissue culture properties, plus 3 representative Type X strains (ARI, RAY, and WTD1), for a total of 19 *T. gondii* strains. Three micrograms of *T. gondii* DNA isolated from each of the 19 strains was sent to NIAID RML for whole-genome sequencing using Illumina HiSeq technology. Fastq reads were reference mapped to the *T. gondii* ME49 assemblage Version 8.2 [70] using BWA 0.7.5a to align the reads to the reference genome and GATK 3.7 in coordination with Picard 1.131, and following best practices *T. gondii* for quality control of mapped reads [77]. Following mapping, the gVCF method of GATK was used to combine SNP calls using stand_call_conf of 30.0, nct of 10, and ploidy of 1 (for haploid genomes) to call 568,592 single nucleotide polymorphism positions across the whole genomes of these strains [77]. The derived VCF formatted SNP file was curated using GATK and VCFTools to produce a tabular file containing only biallelic SNPs with no large insertions or deletions [81]. A custom script was then utilized to convert this SNP file into a fasta file of strain polymorphic positions across the Type X and reference genomes. These SNP fasta files were run in SplitsTree4 using default parameters for BioNJ with 1000 bootstrap support to create a NeighborNet tree [70]. Interconnected reticulation between strains was indicative of recombination between strains, whereas mitotic drift was visualized by divergent branching.

### SNP Density Fingerprint Analysis of Recombinant Progeny

The same tabular, biallelic SNP file used in the NeighborNet analysis was modified using a custom R script based on location mapping to isolate strain-specific polymorphic locations from the tabular VCF. Further R scripts grouped these SNPs into 100 kb windows, which were mapped across the genomes of the sea otter-origin and representative Type X *T. gondii* strains to highlight differences from the reference genome (ME49) [44, 82]. The larger the number of SNPs in a particular window, the more divergent the strain was from the ME49 reference. Distinct haploblocks where genomic recombination had occurred were apparent in areas where haploblock diversity significantly varied from the surrounding regions on the same chromosome.

### PopNet Characterization of Strain Interrelatedness

The tabular, biallelic SNP file created from WGS of Type X and used to derive the SNP density plots and NeighborNet tree was uploaded into PopNet using default parameters to assess the diversity and interrelatedness of strains [38]. Cytoscape was used to visualize the recombination and Markov clustering outputs. Genomes were displayed in circularized format with chromosomes concatenated into a circular genome display. The background of each isolate was painted to match with the group that shared the most common ancestry over the entire genome. Chromosome painting was done in 10 kb increments and the sequence haploblock was painted based on its shared ancestry. For instance, a Type X isolate haploblock that was most closely related to the ancestral Type II strain was painted green to indicate its inheritance. Strains that were more closely related had thicker connecting lines between the circles for each of the *T. gondii* isolates.

### Virulence Gene Identification by QTL

To identify novel virulence alleles, the tabular, biallelic SNP file was utilized to run a quantitative trait locus (QTL) analysis on the Type X and Type II *T. gondii* isolates that were sequenced to WGS resolution [83]. SNP calls were down-selected by a custom java script to include one SNP every 5 kb. This allows the QTL software to analyze the breadth of the WGS data in QTL, which was built to handle marker typing data without the depth inherent in WGS data. Custom scripts coded the SNPs into reference (ME49) versus alternative (Type X’s γ/δ lineage which substituted as the secondary parent) alleles. This down-selected marker dataset was combined with previously determined low-dose inoculum murine virulence data for the same strains. This combined data was then inputted into J/qtl and a one QTL genome scan was run using the default settings for the EM algorithm (maximum likelihood) with 1000 permutations to identify genomic locations that were significantly associated with mouse death due to Type X infection [84].

The QTL calculations identified four genomic regions with a LOD score of greater than 4.0 that were significantly associated with an enhanced risk of mouse death. Within these genomic regions, 450 genes were predicted based on ToxoDB documentation. These potential genes were down-selected based on presence of a signal peptide and/or transmembrane domain, gene expression, polymorphism, and genome-wide CRISPR score for essentiality to identify 32 virulence candidate genes within the Type X strains. Of these, ROP33 was selected for further interrogation based on its high LOD score and similarity to previously identified virulence effector proteins.

## Ethics statements

The animal study protocol LPD 22E was reviewed and approved by the Animal Care and Use Committee of the Intramural Research Program of the National Institute of Allergy and Infectious Diseases, National Institutes of Health. No human studies or identifiable human data are presented in this study. All datasets generated for this study are available upon request to the corresponding author.

## Acknowledgements

We thank Patricia Conrad for critically reading and providing comments on the manuscript. We thank Andrea Packham and Ann Melli for their assistance in culturing and isolation of the *T. gondii* isolates, and Javi Zhang for his help with PopNet analysis. We thank Michael Murray of the Monterey Bay Aquarium and Frances Gulland of the Marine Mammal Center for the submission of additional sea otter cases that yielded *T. gondii* isolates that were not previously characterized. We also thank all volunteers and staff at the California Department of Fish and Wildlife, the Monterey Bay Aquarium, the Marine Mammal Center, and the United States Geological Survey for their efforts to recover stranded sea otters that facilitated the isolation of *T. gondii* strains utilized in this paper. This study used the Office of Cyber Infrastructure and Computational Biology (OCICB) High Performance Computing (HPC) cluster at the National Institute of Allergy and Infectious Diseases (NIAID), Bethesda, MD.

---

**Supplemental Figure 1:**
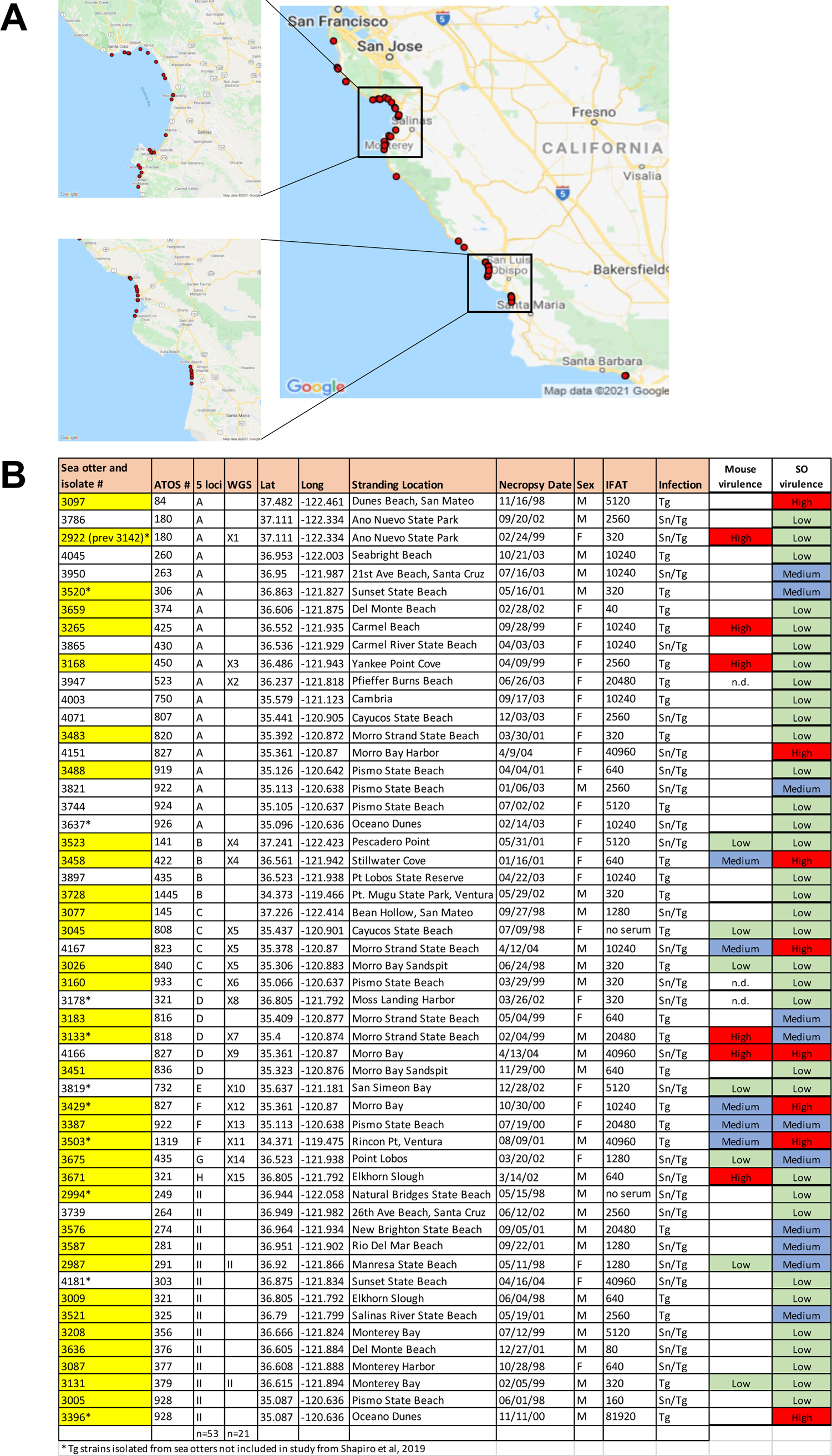
Stranding location, necropsy date, sex, *Toxoplasma* IgG titer (IFAT), and *Sarcocystis neurona* co-infection status for *Toxoplasma gondii* isolates recovered from southern sea otters (1998-2004). A) Coastal California stranding locations for all *Toxoplasma gondii* isolates recovered from southern sea otters (1998-2004). Each stranding location is represented by a red dot. B) *Toxoplasma gondii* isolates included in this study. Isolates in yellow were genotyped previously using PCR-RFLP analyses at 4 loci (*B1, SAG1, SAG2, SAG3*), and isolates 3131, 3133, 3160, 3265 were DNA sequenced at the *GRA6 locus* [25]. WGS analysis further resolved the 21 isolates into 15 distinct Type X genotypes (X1-X15) and two Type II strains. ATOS numbers and stranding locations identify the location of each stranded Sea otter. Isolates were grouped together based on their 5 locus DNA sequence genotype and then by their ATOS number. “Tg” indicates *Toxoplasma gondii,* “Sn” indicates co-infection with *Sarcocystis neurona*.

**Supplemental Table 1:**
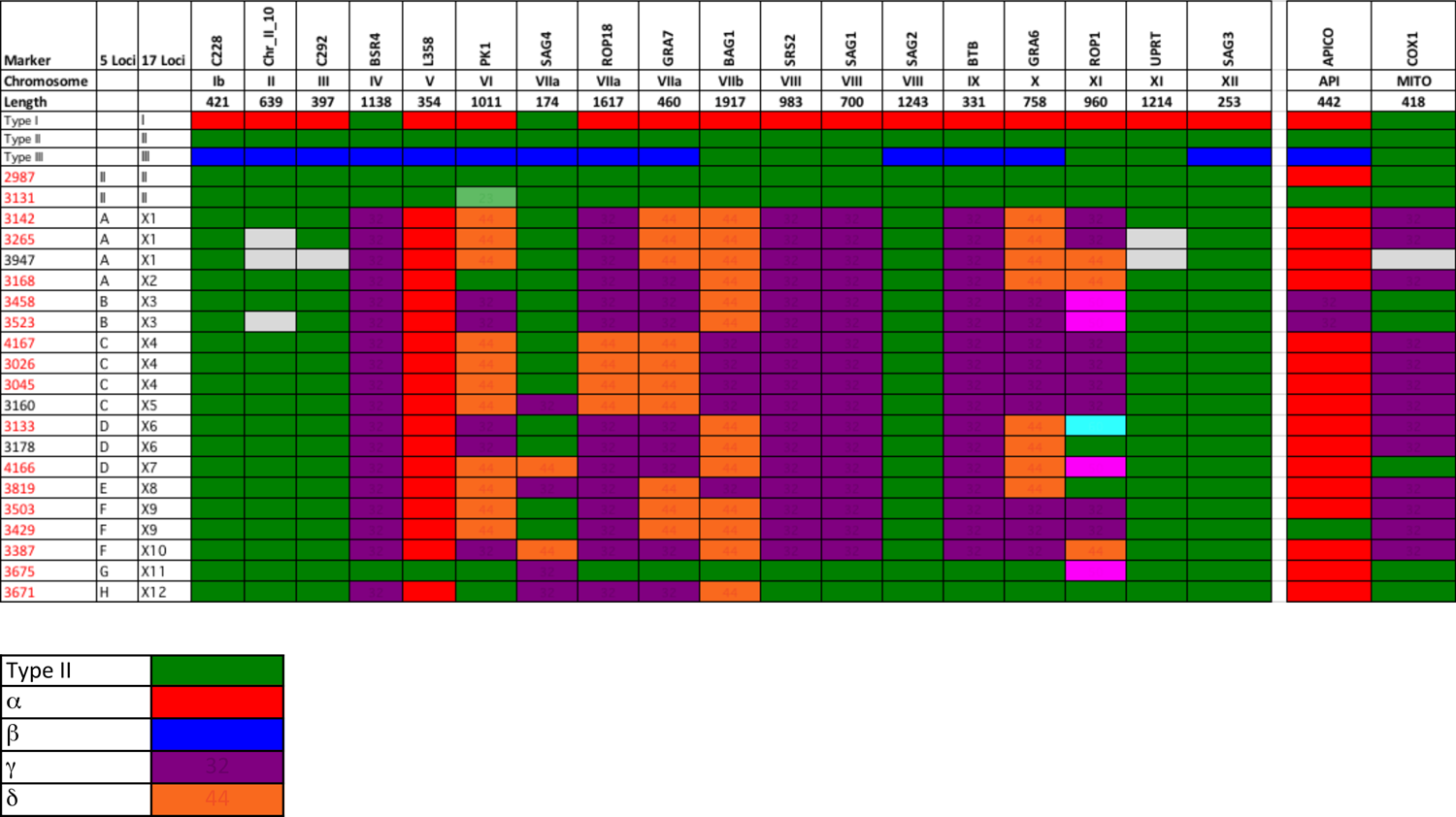
Expanded sequencing marker genotyping of *Toxoplasma gondii* Type X isolates recovered from southern sea otters reveals chromosomal segregation and recombination within chromosomes Twenty-one isolates were characterized at 17 nuclear markers (ROP1 was excluded as it is a microsatellite marker, and prone to elevated mutation rates) clade into 12 distinct Type X genotypes (X1-X12). Isolate numbers in red were sequenced at whole-genome (WGS) resolution. Types I, II, III, γ and δ lineage alleles, as determined by phylogenetic comparisons, are colored red, green, blue, purple, and orange, respectively. Shades of colors represent genetic drift below the 60% bootstrap delineation from the canonical allele. White represents uninformative or incomplete DNA sequencing results.

**Supplemental Table 2:**
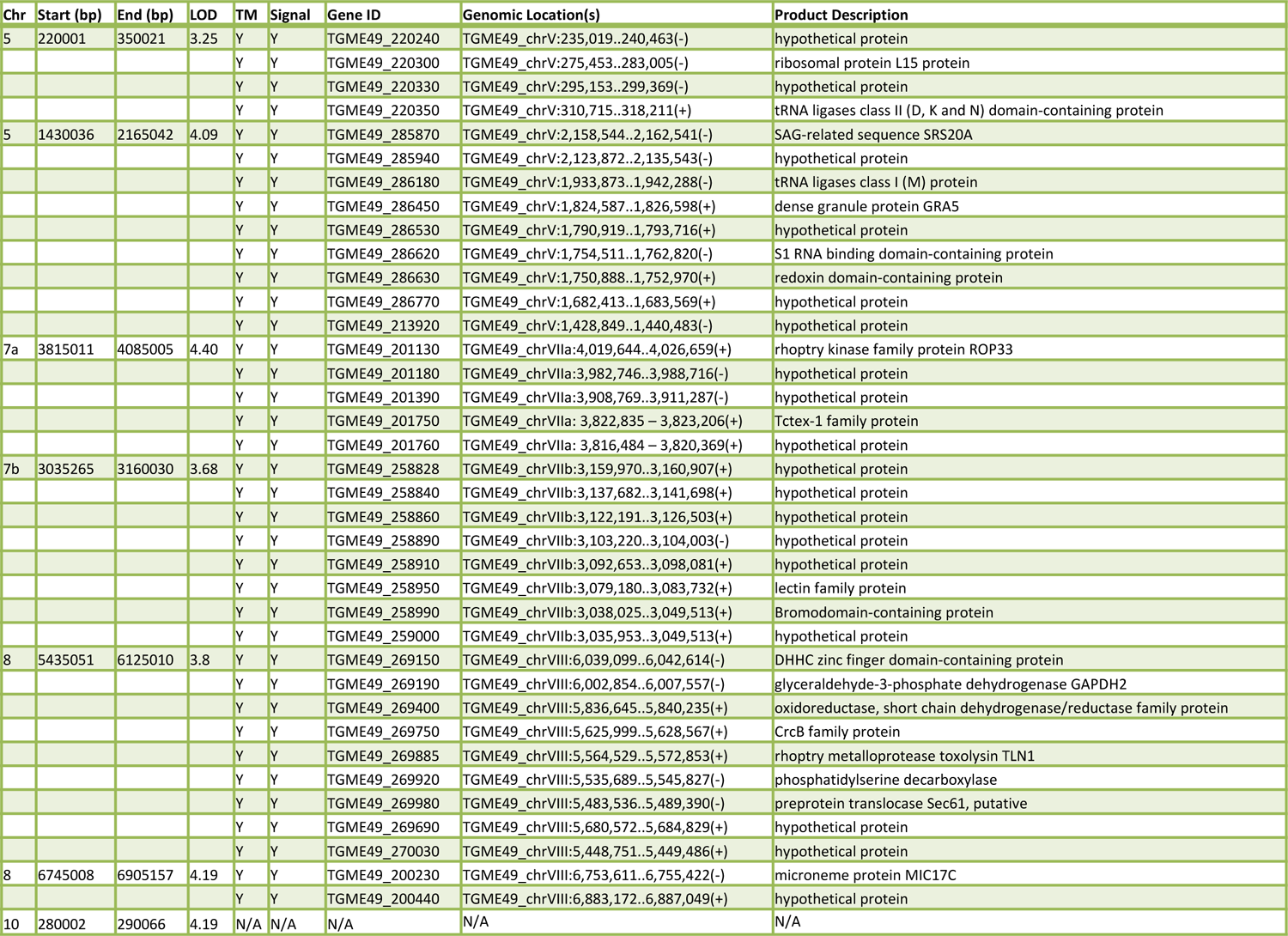
Potential virulence candidate genes derived from the QTL analysis Chromosome peaks with LOD scores over 3 as shown in Figure 3B were interrogated via ToxoDB to select genes in these regions which have predicted signal sequences (Signal) and transmembrane (TM) domains in *Toxoplasma.* Genes are listed with gene ID, genomic location, and predicted protein function.

## References

1. Vijaykrishna D, Smith GJ, Pybus OG, Zhu H, Bhatt S, Poon LL, et al. Long-term evolution and transmission dynamics of swine influenza A virus. Nature. 2011;473(7348):519-22. doi: 10.1038/nature10004. PubMed PMID: 21614079.

2. Bellanger X, Payot S, Leblond-Bourget N, Guedon G. Conjugative and mobilizable genomic islands in bacteria: evolution and diversity. FEMS Microbiol Rev. 2014;38(4):720–60. doi: 10.1111/1574-6976.12058. PubMed PMID: 24372381.

3. Schneider G, Dobrindt U, Middendorf B, Hochhut B, Szijarto V, Emody L, et al. Mobilisation and remobilisation of a large archetypal pathogenicity island of uropathogenic Escherichia coli in vitro support the role of conjugation for horizontal transfer of genomic islands. BMC Microbiol. 2011;11:210. doi: 10.1186/1471-2180-11-210. PubMed PMID: 21943043; PubMed Central PMCID: PMC3202238.

4. Wardal E, Markowska K, Zabicka D, Wroblewska M, Giemza M, Mik E, et al. Molecular analysis of vanA outbreak of Enterococcus faecium in two Warsaw hospitals: the importance of mobile genetic elements. Biomed Res Int. 2014;2014:575367. doi: 10.1155/2014/575367. PubMed PMID: 25003118; PubMed Central PMCID: PMC4070583.

5. English ED, Adomako-Ankomah Y, Boyle JP. Secreted effectors in Toxoplasma gondii and related species: determinants of host range and pathogenesis? Parasite Immunol. 2015;37(3):127–40. doi: 10.1111/pim.12166. PubMed PMID: 25655311.

6. Fraser JA, Giles SS, Wenink EC, Geunes-Boyer SG, Wright JR, Diezmann S, et al. Same-sex mating and the origin of the Vancouver Island Cryptococcus gattii outbreak. Nature. 2005;437(7063):1360-4. Epub 2005/10/14. doi: 10.1038/nature04220. PubMed PMID: 16222245.

7. Grigg ME, Bonnefoy S, Hehl AB, Suzuki Y, Boothroyd JC. Success and virulence in toxoplasma as the result of sexual recombination between two distinct ancestries. Science. 2001;294(5540):161-5. doi: DOI 10.1126/science.1061888. PubMed PMID: WOS:000171448800052.

8. Boothroyd JC, Grigg ME. Population biology of Toxoplasma gondii and its relevance to human infection: do different strains cause different disease? Curr Opin Microbiol. 5. England 2002. p. 438-42.

9. Boyer K, Hill D, Mui E, Wroblewski K, Karrison T, Dubey JP, et al. Unrecognized ingestion of Toxoplasma gondii oocysts leads to congenital toxoplasmosis and causes epidemics in North America. Clin Infect Dis. 2011;53(11):1081–9. Epub 2011/10/25. doi: 10.1093/cid/cir667. PubMed PMID: 22021924; PubMed Central PMCID: PMCPMC3246875.

10. Behnke MS, Khan A, Wootton JC, Dubey JP, Tang K, Sibley LD. Virulence differences in Toxoplasma mediated by amplification of a family of polymorphic pseudokinases. Proc Natl Acad Sci U S A. 2011;108(23):9631–6. doi: 10.1073/pnas.1015338108. PubMed PMID: 21586633; PubMed Central PMCID: PMC3111276.

11. Boyle JP, Rajasekar B, Saeij JP, Ajioka JW, Berriman M, Paulsen I, et al. Just one cross appears capable of dramatically altering the population biology of a eukaryotic pathogen like Toxoplasma gondii. Proc Natl Acad Sci U S A. 2006;103(27):10514–9. doi: 10.1073/pnas.0510319103. PubMed PMID: 16801557; PubMed Central PMCID: PMC1502489.

12. Wendte JM, Miller MA, Lambourn DM, Magargal SL, Jessup DA, Grigg ME. Self-mating in the definitive host potentiates clonal outbreaks of the apicomplexan parasites Sarcocystis neurona and Toxoplasma gondii. PLoS Genet. 2010;6(12):e1001261. doi: 10.1371/journal.pgen.1001261. PubMed PMID: 21203443; PubMed Central PMCID: PMC3009688.

13. Boothroyd JC. Expansion of host range as a driving force in the evolution of Toxoplasma. Memorias Do Instituto Oswaldo Cruz. 2009;104(2):179–84. PubMed PMID: WOS:000267051500009.

14. Howe DK, Sibley LD. Toxoplasma gondii comprises three clonal lineages: correlation of parasite genotype with human disease. J Infect Dis. 1995;172(6):1561–6. PubMed PMID: 7594717.

15. Lorenzi H, Khan A, Behnke MS, Namasivayam S, Swapna LS, Hadjithomas M, et al. Local admixture of amplified and diversified secreted pathogenesis determinants shapes mosaic Toxoplasma gondii genomes. Nat Commun. 2016;7:10147. doi: 10.1038/ncomms10147. PubMed PMID: 26738725; PubMed Central PMCID: PMCPMC4729833.

16. Sibley LD, Boothroyd JC. Virulent strains of Toxoplasma gondii comprise a single clonal lineage. Nature. 1992;359(6390):82-5. doi: 10.1038/359082a0. PubMed PMID: 1355855.

17. Lilue J, Muller UB, Steinfeldt T, Howard JC. Reciprocal virulence and resistance polymorphism in the relationship between Toxoplasma gondii and the house mouse. Elife. 2013;2:e01298. doi: 10.7554/eLife.01298. PubMed PMID: 24175088; PubMed Central PMCID: PMC3810784.

18. Steinfeldt T, Konen-Waisman S, Tong L, Pawlowski N, Lamkemeyer T, Sibley LD, et al. Phosphorylation of mouse immunity-related GTPase (IRG) resistance proteins is an evasion strategy for virulent Toxoplasma gondii. PLoS Biol. 2010;8(12):e1000576. Epub 2011/01/05. doi: 10.1371/journal.pbio.1000576. PubMed PMID: 21203588; PubMed Central PMCID: PMCPMC3006384.

19. Cavailles P, Sergent V, Bisanz C, Papapietro O, Colacios C, Mas M, et al. The rat Toxo1 locus directs toxoplasmosis outcome and controls parasite proliferation and spreading by macrophage-dependent mechanisms. Proc Natl Acad Sci U S A. 2006;103(3):744–9. Epub 2006/01/13. doi: 10.1073/pnas.0506643103. PubMed PMID: 16407112; PubMed Central PMCID: PMCPMC1334643.

20. Cirelli KM, Gorfu G, Hassan MA, Printz M, Crown D, Leppla SH, et al. Inflammasome sensor NLRP1 controls rat macrophage susceptibility to Toxoplasma gondii. PLoS Pathog. 2014;10(3):e1003927. Epub 2014/03/15. doi: 10.1371/journal.ppat.1003927. PubMed PMID: 24626226; PubMed Central PMCID: PMCPMC3953412.

21. Ewald SE, Chavarria-Smith J, Boothroyd JC. NLRP1 is an inflammasome sensor for Toxoplasma gondii. Infect Immun. 2014;82(1):460–8. Epub 2013/11/13. doi: 10.1128/IAI.01170-13. PubMed PMID: 24218483; PubMed Central PMCID: PMCPMC3911858.

22. Gorfu G, Cirelli KM, Melo MB, Mayer-Barber K, Crown D, Koller BH, et al. Dual role for inflammasome sensors NLRP1 and NLRP3 in murine resistance to Toxoplasma gondii. MBio. 2014;5(1). Epub 2014/02/20. doi: 10.1128/mBio.01117-13. PubMed PMID: 24549849; PubMed Central PMCID: PMCPMC3944820.

23. Dubey JP, Shen SK, Kwok OC, Frenkel JK. Infection and immunity with the RH strain of Toxoplasma gondii in rats and mice. J Parasitol. 1999;85(4):657–62. PubMed PMID: 10461945.

24. Conrad PA, Miller MA, Kreuder C, James ER, Mazet J, Dabritz H, et al. Transmission of Toxoplasma: clues from the study of sea otters as sentinels of Toxoplasma gondii flow into the marine environment. Int J Parasitol. 2005;35(11-12):1155–68. Epub 2005/09/15. doi: 10.1016/j.ijpara.2005.07.002. PubMed PMID: 16157341.

25. Miller MA, Grigg ME, Kreuder C, James ER, Melli AC, Crosbie PR, et al. An unusual genotype of Toxoplasma gondii is common in California sea otters (Enhydra lutris nereis) and is a cause of mortality. Int J Parasitol. 2004;34(3):275–84. doi: 10.1016/j.ijpara.2003.12.008. PubMed PMID: 15003489.

26. Shapiro K, VanWormer E, Packham A, Dodd E, Conrad PA, Miller M. Type X strains of Toxoplasma gondii are virulent for southern sea otters (Enhydra lutris nereis) and present in felids from nearby watersheds. Proc Biol Sci. 2019;286(1909):20191334. Epub 2019/08/23. doi: 10.1098/rspb.2019.1334. PubMed PMID: 31431162; PubMed Central PMCID: PMCPMC6732395.

27. Khan A, Dubey JP, Su C, Ajioka JW, Rosenthal BM, Sibley LD. Genetic analyses of atypical Toxoplasma gondii strains reveal a fourth clonal lineage in North America. Int J Parasitol. 2011;41(6):645–55. doi: 10.1016/j.ijpara.2011.01.005. PubMed PMID: 21320505; PubMed Central PMCID: PMC3081397.

28. Dubey JP, Velmurugan GV, Rajendran C, Yabsley MJ, Thomas NJ, Beckmen KB, et al. Genetic characterisation of Toxoplasma gondii in wildlife from North America revealed widespread and high prevalence of the fourth clonal type. Int J Parasitol. 2011;41(11):1139–47. doi: 10.1016/j.ijpara.2011.06.005. PubMed PMID: 21802422.

29. VanWormer E, Miller MA, Conrad PA, Grigg ME, Rejmanek D, Carpenter TE, et al. Using molecular epidemiology to track Toxoplasma gondii from terrestrial carnivores to marine hosts: implications for public health and conservation. PLoS Negl Trop Dis. 2014;8(5):e2852. doi: 10.1371/journal.pntd.0002852. PubMed PMID: 24874796; PubMed Central PMCID: PMC4038486.

30. Gibson AK, Raverty S, Lambourn DM, Huggins J, Magargal SL, Grigg ME. Polyparasitism is associated with increased disease severity in Toxoplasma gondii-infected marine sentinel species. PLoS Negl Trop Dis. 2011;5(5):e1142. doi: 10.1371/journal.pntd.0001142. PubMed PMID: 21629726; PubMed Central PMCID: PMC3101184.

31. Kreuder C, Miller MA, Jessup DA, Lowenstine LJ, Harris MD, Ames JA, et al. Patterns of mortality in southern sea otters (Enhydra lutris nereis) from 1998-2001. J Wildl Dis. 2003;39(3):495–509. doi: 10.7589/0090-3558-39.3.495. PubMed PMID: 14567210.

32. Sundar N, Cole RA, Thomas NJ, Majumdar D, Dubey JP, Su C. Genetic diversity among sea otter isolates of Toxoplasma gondii. Vet Parasitol. 2008;151(2-4):125–32. doi: 10.1016/j.vetpar.2007.11.012. PubMed PMID: 18155841.

33. Beraki T, Hu X, Broncel M, Young JC, O’Shaughnessy WJ, Borek D, et al. Divergent kinase regulates membrane ultrastructure of the Toxoplasma parasitophorous vacuole. Proc Natl Acad Sci U S A. 2019;116(13):6361–70. Epub 2019/03/10. doi: 10.1073/pnas.1816161116. PubMed PMID: 30850550; PubMed Central PMCID: PMCPMC6442604.

34. Miller MA, Miller WA, Conrad PA, James ER, Melli AC, Leutenegger CM, et al. Type X Toxoplasma gondii in a wild mussel and terrestrial carnivores from coastal California: new linkages between terrestrial mammals, runoff and toxoplasmosis of sea otters. Int J Parasitol. 2008;38(11):1319–28. doi: 10.1016/j.ijpara.2008.02.005. PubMed PMID: 18452923.

35. Su C, Khan A, Zhou P, Majumdar D, Ajzenberg D, Darde ML, et al. Globally diverse Toxoplasma gondii isolates comprise six major clades originating from a small number of distinct ancestral lineages. Proc Natl Acad Sci U S A. 2012;109(15):5844–9. doi: 10.1073/pnas.1203190109. PubMed PMID: 22431627; PubMed Central PMCID: PMCPMC3326454.

36. Khan A, Miller N, Roos DS, Dubey JP, Ajzenberg D, Darde ML, et al. A monomorphic haplotype of chromosome Ia is associated with widespread success in clonal and nonclonal populations of Toxoplasma gondii. MBio. 2011;2(6):e00228–11. doi: 10.1128/mBio.00228-11. PubMed PMID: 22068979; PubMed Central PMCID: PMC3215432.

37. Zhang J, Khan A, Kennard A, Grigg ME, Parkinson J. PopNet: A Markov Clustering Approach to Study Population Genetic Structure. Mol Biol Evol. 2017;34(7):1799–811. Epub 2017/04/07. doi: 10.1093/molbev/msx110. PubMed PMID: 28383661; PubMed Central PMCID: PMCPMC5850731.

38. Sidik SM, Huet D, Ganesan SM, Huynh MH, Wang T, Nasamu AS, et al. A Genome-wide CRISPR Screen in Toxoplasma Identifies Essential Apicomplexan Genes. Cell. 2016;166(6):1423–35 e12. Epub 2016/09/07. doi: 10.1016/j.cell.2016.08.019. PubMed PMID: 27594426; PubMed Central PMCID: PMCPMC5017925.

39. Etheridge RD, Alaganan A, Tang K, Lou HJ, Turk BE, Sibley LD. The Toxoplasma pseudokinase ROP5 forms complexes with ROP18 and ROP17 kinases that synergize to control acute virulence in mice. Cell Host Microbe. 2014;15(5):537–50. Epub 2014/05/17. doi: 10.1016/j.chom.2014.04.002. PubMed PMID: 24832449; PubMed Central PMCID: PMCPMC4086214.

40. Khan A, Fux B, Su C, Dubey JP, Darde ML, Ajioka JW, et al. Recent transcontinental sweep of Toxoplasma gondii driven by a single monomorphic chromosome. Proc Natl Acad Sci U S A. 2007;104(37):14872–7. Epub 2007/09/07. doi: 10.1073/pnas.0702356104. PubMed PMID: 17804804; PubMed Central PMCID: PMCPMC1965483.

41. Grigg ME, Sundar N. Sexual recombination punctuated by outbreaks and clonal expansions predicts Toxoplasma gondii population genetics. Int J Parasitol. 2009;39(8):925–33. doi: 10.1016/j.ijpara.2009.02.005. PubMed PMID: 19217909; PubMed Central PMCID: PMC2713429.

42. Minot S, Melo MB, Li F, Lu D, Niedelman W, Levine SS, et al. Admixture and recombination among Toxoplasma gondii lineages explain global genome diversity. Proceedings of the National Academy of Sciences of the United States of America. 2012;109(33):13458–63. doi: DOI 10.1073/pnas.1117047109. PubMed PMID: WOS:000307807000067; PubMed Central PMCID: PMCPMC3421188.

43. Khan A, Shaik JS, Behnke M, Wang QL, Dubey JP, Lorenzi HA, et al. NextGen sequencing reveals short double crossovers contribute disproportionately to genetic diversity in Toxoplasma gondii. Bmc Genomics. 2014;15. doi: Artn 1168 10.1186/1471-2164-15-1168. PubMed PMID: WOS:000349049000003.

44. Rajendran C, Su C, Dubey JP. Molecular genotyping of Toxoplasma gondii from Central and South America revealed high diversity within and between populations. Infect Genet Evol. 2012;12(2):359–68. Epub 2012/01/10. doi: 10.1016/j.meegid.2011.12.010. PubMed PMID: 22226702.

45. Saeij JP, Boyle JP, Coller S, Taylor S, Sibley LD, Brooke-Powell ET, et al. Polymorphic secreted kinases are key virulence factors in toxoplasmosis. Science. 2006;314(5806):1780-3. doi: 10.1126/science.1133690. PubMed PMID: 17170306; PubMed Central PMCID: PMC2646183.

46. Saeij JP, Coller S, Boyle JP, Jerome ME, White MW, Boothroyd JC. Toxoplasma co-opts host gene expression by injection of a polymorphic kinase homologue. Nature. 2007;445(7125):324-7. doi: 10.1038/nature05395. PubMed PMID: 17183270; PubMed Central PMCID: PMC2637441.

47. Taylor S, Barragan A, Su C, Fux B, Fentress SJ, Tang K, et al. A secreted serine-threonine kinase determines virulence in the eukaryotic pathogen Toxoplasma gondii. Science. 2006;314(5806):1776-80. doi: 10.1126/science.1133643. PubMed PMID: 17170305.

48. Ferguson D. Toxoplasma gondii and sex: essential or optional extra? Trends Parasitol. 2002;18(8):351. Epub 2002/10/16. PubMed PMID: 12377284.

49. Fleckenstein MC, Reese ML, Konen-Waisman S, Boothroyd JC, Howard JC, Steinfeldt T. A Toxoplasma gondii pseudokinase inhibits host IRG resistance proteins. PLoS Biol. 2012;10(7):e1001358. doi: 10.1371/journal.pbio.1001358. PubMed PMID: 22802726; PubMed Central PMCID: PMC3393671.

50. Hunn JP, Feng CG, Sher A, Howard JC. The immunity-related GTPases in mammals: a fast-evolving cell-autonomous resistance system against intracellular pathogens. Mamm Genome. 2011;22(1-2):43–54. doi: 10.1007/s00335-010-9293-3. PubMed PMID: 21052678; PubMed Central PMCID: PMC3438224.

51. Yamamoto M, Okuyama M, Ma JS, Kimura T, Kamiyama N, Saiga H, et al. A cluster of interferon-gamma-inducible p65 GTPases plays a critical role in host defense against Toxoplasma gondii. Immunity. 2012;37(2):302–13. doi: 10.1016/j.immuni.2012.06.009. PubMed PMID: 22795875.

52. Gazzinelli RT, Mendonca-Neto R, Lilue J, Howard J, Sher A. Innate Resistance against Toxoplasma gondii: An Evolutionary Tale of Mice, Cats, and Men. Cell Host Microbe. 2014;15(2):132–8. Epub 2014/02/18. doi: 10.1016/j.chom.2014.01.004. PubMed PMID: 24528860.

53. Koblansky AA, Jankovic D, Oh H, Hieny S, Sungnak W, Mathur R, et al. Recognition of profilin by Toll-like receptor 12 is critical for host resistance to Toxoplasma gondii. Immunity. 2013;38(1):119–30. doi: 10.1016/j.immuni.2012.09.016. PubMed PMID: 23246311; PubMed Central PMCID: PMC3601573.

54. Mackinnon MJ, Read AF. Virulence in malaria: an evolutionary viewpoint. Philos Trans R Soc Lond B Biol Sci. 2004;359(1446):965-86. doi: 10.1098/rstb.2003.1414. PubMed PMID: 15306410; PubMed Central PMCID: PMC1693375.

55. Ariey F, Witkowski B, Amaratunga C, Beghain J, Langlois AC, Khim N, et al. A molecular marker of artemisinin-resistant Plasmodium falciparum malaria. Nature. 2014;505(7481):50-5. Epub 2013/12/20. doi: 10.1038/nature12876. PubMed PMID: 24352242.

56. Bopp SE, Manary MJ, Bright AT, Johnston GL, Dharia NV, Luna FL, et al. Mitotic evolution of Plasmodium falciparum shows a stable core genome but recombination in antigen families. PLoS Genet. 2013;9(2):e1003293. doi: 10.1371/journal.pgen.1003293. PubMed PMID: 23408914; PubMed Central PMCID: PMC3567157.

57. Miotto O, Almagro-Garcia J, Manske M, Macinnis B, Campino S, Rockett KA, et al. Multiple populations of artemisinin-resistant Plasmodium falciparum in Cambodia. Nat Genet. 2013;45(6):648–55. Epub 2013/04/30. doi: 10.1038/ng.2624. PubMed PMID: 23624527; PubMed Central PMCID: PMC3807790.

58. Arisue N, Hashimoto T. Phylogeny and evolution of apicoplasts and apicomplexan parasites. Parasitol Int. 2014. doi: 10.1016/j.parint.2014.10.005. PubMed PMID: 25451217.

59. Barbosa L, Johnson CK, Lambourn DM, Gibson AK, Haman KH, Huggins JL, et al. A novel Sarcocystis neurona genotype XIII is associated with severe encephalitis in an unexpectedly broad range of marine mammals from the northeastern Pacific Ocean. Int J Parasitol. 2015. doi: 10.1016/j.ijpara.2015.02.013. PubMed PMID: 25997588.

60. Dubey JP, Graham DH, da Silva DS, Lehmann T, Bahia-Oliveira LM. Toxoplasma gondii isolates of free-ranging chickens from Rio de Janeiro, Brazil: mouse mortality, genotype, and oocyst shedding by cats. J Parasitol. 2003;89(4):851–3. Epub 2003/10/10. doi: 10.1645/GE-60R. PubMed PMID: 14533703.

61. Wendte JM, Gibson AK, Grigg ME. Population genetics of Toxoplasma gondii: new perspectives from parasite genotypes in wildlife. Vet Parasitol. 2011;182(1):96–111. doi: 10.1016/j.vetpar.2011.07.018. PubMed PMID: 21824730; PubMed Central PMCID: PMC3430134.

62. Labruyere E, Lingnau M, Mercier C, Sibley LD. Differential membrane targeting of the secretory proteins GRA4 and GRA6 within the parasitophorous vacuole formed by Toxoplasma gondii. Mol Biochem Parasitol. 1999;102(2):311–24. Epub 1999/09/25. doi: 10.1016/s0166-6851(99)00092-4. PubMed PMID: 10498186.

63. Mercier C, Howe DK, Mordue D, Lingnau M, Sibley LD. Targeted disruption of the GRA2 locus in Toxoplasma gondii decreases acute virulence in mice. Infect Immun. 1998;66(9):4176–82. Epub 1998/08/26. PubMed PMID: 9712765; PubMed Central PMCID: PMCPMC108503.

64. Wang JL, Li TT, Elsheikha HM, Chen K, Zhu WN, Yue DM, et al. Functional Characterization of Rhoptry Kinome in the Virulent Toxoplasma gondii RH Strain. Front Microbiol. 2017;8:84. Epub 2017/02/09. doi: 10.3389/fmicb.2017.00084. PubMed PMID: 28174572; PubMed Central PMCID: PMCPMC5258691.

65. Miller MA, Gardner IA, Packham A, Mazet JK, Hanni KD, Jessup D, et al. Evaluation of an Indirect Fluorescent Antibody Test (Ifat) for Demonstration of Antibodies to Toxoplasma Gondii in the Sea Otter (Enhydra Lutris). Journal of Parasitology. 2002;88(3):594–9. doi: 10.1645/0022-3395(2002)088[0594:eoaifa]2.0.co;2.

66. Pszenny V, Angel SO, Duschak VG, Paulino M, Ledesma B, Yabo MI, et al. Molecular cloning, sequencing and expression of a serine proteinase inhibitor gene from Toxoplasma gondii. Mol Biochem Parasitol. 2000;107(2):241–9. PubMed PMID: 10779600.

67. Su C, Zhang X, Dubey JP. Genotyping of Toxoplasma gondii by multilocus PCR-RFLP markers: a high resolution and simple method for identification of parasites. Int J Parasitol. 2006;36(7):841–8. doi: 10.1016/j.ijpara.2006.03.003. PubMed PMID: 16643922.

68. Fazaeli A, Carter PE, Darde ML, Pennington TH. Molecular typing of Toxoplasma gondii strains by GRA6 gene sequence analysis. Int J Parasitol. 2000;30(5):637–42. PubMed PMID: 10779578.

69. Howe DK, Sibley LD. Toxoplasma gondii: analysis of different laboratory stocks of the RH strain reveals genetic heterogeneity. Exp Parasitol. 1994;78(2):242–5. doi: 10.1006/expr.1994.1024. PubMed PMID: 7907030.

70. Gajria B, Bahl A, Brestelli J, Dommer J, Fischer S, Gao X, et al. ToxoDB: an integrated Toxoplasma gondii database resource. Nucleic Acids Res. 2008;36(Database issue):D553-6. doi: 10.1093/nar/gkm981. PubMed PMID: 18003657; PubMed Central PMCID: PMC2238934.

71. Donald RG, Carter D, Ullman B, Roos DS. Insertional tagging, cloning, and expression of the Toxoplasma gondii hypoxanthine-xanthine-guanine phosphoribosyltransferase gene. Use as a selectable marker for stable transformation. J Biol Chem. 1996;271(24):14010–9. Epub 1996/06/14. doi: 10.1074/jbc.271.24.14010. PubMed PMID: 8662859.

72. Fletcher S. Indirect Fluorescent Antibody Technique in the Serology of Toxoplasma Gondii. J Clin Pathol. 1965;18:193–9. Epub 1965/03/01. PubMed PMID: 14276154; PubMed Central PMCID: PMC472865.

73. Larkin MA, Blackshields G, Brown NP, Chenna R, McGettigan PA, McWilliam H, et al. Clustal W and Clustal X version 2.0. Bioinformatics. 2007;23(21):2947–8. doi: 10.1093/bioinformatics/btm404. PubMed PMID: 17846036.

74. Tamura K, Stecher G, Peterson D, Filipski A, Kumar S. MEGA6: Molecular Evolutionary Genetics Analysis version 6.0. Mol Biol Evol. 2013;30(12):2725–9. doi: 10.1093/molbev/mst197. PubMed PMID: 24132122; PubMed Central PMCID: PMC3840312.

75. Huson DH, Bryant D. Application of phylogenetic networks in evolutionary studies. Mol Biol Evol. 2006;23(2):254–67. doi: 10.1093/molbev/msj030. PubMed PMID: 16221896.

76. Feil EJ, Li BC, Aanensen DM, Hanage WP, Spratt BG. eBURST: Inferring Patterns of Evolutionary Descent among Clusters of Related Bacterial Genotypes from Multilocus Sequence Typing Data. Journal of Bacteriology. 2004;186(5):1518–30. doi: 10.1128/jb.186.5.1518-1530.2004.

77. Van der Auwera GA, Carneiro MO, Hartl C, Poplin R, Del Angel G, Levy-Moonshine A, et al. From FastQ data to high confidence variant calls: the Genome Analysis Toolkit best practices pipeline. Curr Protoc Bioinformatics. 2013;11(1110):11 0 1-0 33. doi: 10.1002/0471250953.bi1110s43. PubMed PMID: 25431634; PubMed Central PMCID: PMC4243306.

